# Futile wound healing drives mesenchymal-like cell phenotypes in human glioblastoma

**DOI:** 10.1101/2023.09.01.555882

**Authors:** Alejandro Mossi Albiach, Jokubas Janusauskas, Jesper Kjaer Jacobsen, Ivana Kapustová, Razieh Karamzadeh, Egle Kvedaraite, Lijuan Hu, Marina C. M. Franck, Camiel Mannens, Simone Codeluppi, Johannes B. Munting, Lars E. Borm, Alia Shamikh, Peter Lönnerberg, Kimberly A. Siletti, Oscar Persson, Sten Linnarsson

## Abstract

Glioblastoma is the deadliest brain cancer, characterized by large cellular diversity whose complexity and organizing principles are only starting to be uncovered. Both neurodevelopment-like and mesenchymal-like cell states have been described in glioblastoma^1–8^, with the latter being strongly implicated in malignancy and disease progression^8–11^. However, the nature of these mesenchymal-like cell states remains unresolved. Here, we performed deep single-cell RNA sequencing of rare glioblastoma cases where tissue could be sampled from tumor core to macroscopically normal cortex. We discovered that previously defined mesenchymal-like tumor cell states instead represented a wound response that was shared across both malignant and non-malignant cell types and was spatially confined to the tumor bulk. Using glioblastoma organoids, we showed that the wound response transcriptional state could be reversibly induced *in vitro* by hypoxia and human plasma. We used multiplex single-molecule spatial transcriptomics^12^ on a large patient cohort to show that the activation of wound response states was associated with hypoxia, and organized by distance to perivascular niches. Our findings help reconceptualize the cellular landscape of glioblastoma, wherein a reactive wound-response tissue state shared by all cells in the tumor bulk is superimposed on a fundamentally neurodevelopmental and glial tumor.

## MAIN

Glioblastoma (GBM) is the most common malignant brain cancer in adults with a median survival of less than a year^13^. While heterogeneous, these tumors nevertheless show phenotypical convergence to common cell states, oftentimes resembling neurodevelopmental lineages and commonly described as astrocyte (AC)-like, oligodendrocyte precursor cell (OPC)-like, neural progenitor cell (NPC)-like, and mesenchymal (MES)-like^1–8^. MES-like tumor cells have been associated with poor prognosis and relapse^8,9^, invasiveness^10^ and resistance to therapy^10,11^. This clinical importance has sparked investigations into the nature of the MES-like state^14–16^ as well as efforts to identify factors that endogenously induce the MES-like phenotype or that pharmacologically block it, as a potential therapy^17–19^. Tumor-associated macrophages (TAMs) have also been shown to induce MES-like cell states^20^, and attempts have been made to target TAMs therapeutically in order to hinder tumor growth^21–23^. However, in mouse models, depletion of TAMs does not slow tumor growth^24^ and can even promote it^25^. Taken together, pinpointing the causal roles of MES-like tumor cells remains a challenging yet central issue in GBM research.

Moreover, GBM surgery follows the principle of maximal safe resection of the MRI contrast-enhancing component, inevitably leaving behind infiltrating tumor cells in the tumor periphery that ultimately cause relapse^26^. This also defines tissue availability, meaning that nearly all previous studies have focused on the contrast-enhancing tumor part (although notable exceptions exist^27,28^) and that our understanding of tumor cell states beyond the tumor bulk is likewise limited. Therefore, to elucidate the role and function of MES-like cell states in GBM, here we took advantage of rare temporal lobectomies, where tissue sampling beyond the tumor margin and into macroscopically unaffected cortex was possible. We performed deep single-cell RNA sequencing on a couple of multi-region GBM cases – sampling more than 100,000 cells each – and spatial transcriptomics using 888-plex Enhanced ELectric single-molecule Fluorescence *in situ* Hybridization (EEL-FISH)^12^ on multi-region samples as well as standard resection samples for a total of 41 tissue samples from 27 gliomas: 25 GBMs and two cases of oligodendroglioma (IDH-mutated, 1p19q co-deleted).

### Mesenchymal-like gene expression is regulated spatially across cell types

To understand the large-scale spatial distribution of MES-like tumor cells, we first examined a rare case of GBM (sample SL040), where location of the tumor required surgical resection far outside of the MRI contrast-enhancing zone. Samples were annotated based on 5-aminolevulinic acid (5-ALA, Gliolan) fluorescence (Methods, **Fig. 1a, Extended Data Fig. 1a**); wherein high fluorescence largely corresponds to the contrast-enhancing zone^29^. We collected tissue from five nested zones: the necrotic core (central tumor mass, low or no fluorescence), high-fluorescence zone (representing the contrast-enhancing zone), low-fluorescence zone (immediately surrounding the high-fluorescence zone), tumor periphery and the surrounding cortex (**Fig. 1a**, **Extended Data Fig. 1b**). Multiple samples were taken in different directions (anterior, posterior, superior, inferior), for a total of 12 sampling locations (Methods**, Extended Data Fig. 1b**).

**Figure 1.**
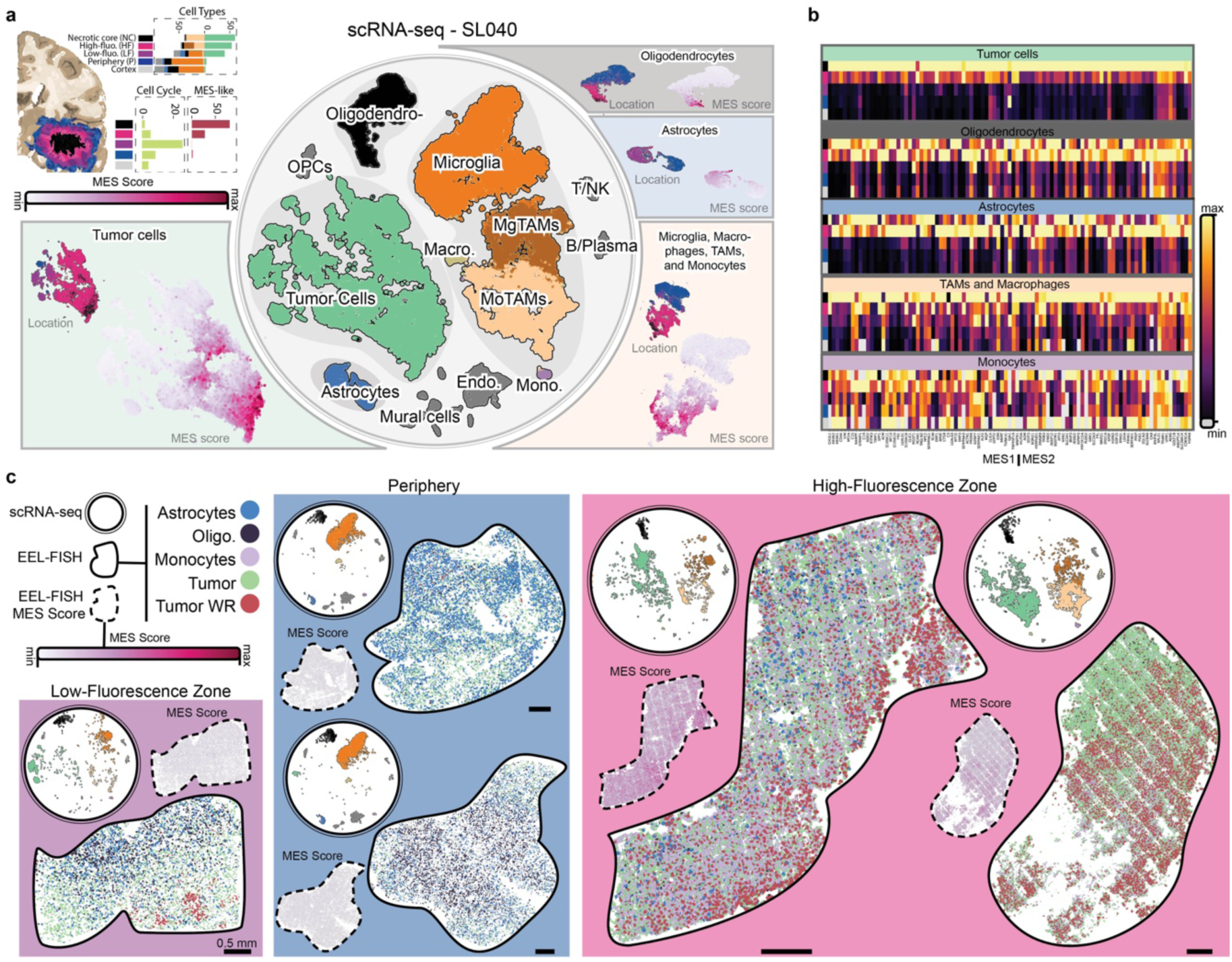
scRNA-seq and EEL-FISH characterization of a multi-region GBM sample. **a**, Top-left: schematic illustration of the organization of the fluorescence zones in GBM as part of a coronal brain section: necrotic core, high-fluorescence zone, low-fluorescence zone, periphery and cortex; cell cycle score and fraction of MES-tumor cells across different tumor zones. Center: t-distributed stochastic neighbor embedding (t-SNE) representation of scRNA-seq data from multi-region sample SL040. Surrounding the center: plots of regional distribution and MES score for tumor cells, astrocytes, oligodendrocytes, monocytes, microglia, macrophages, TAMs and other major cell types. **b,** MES-like gene expression across cell types and fluorescence zones. The colormap shows linear gene expression truncated at the 99.5% percentile, from the minimum to the maximum value per gene, with zeros shown in gray. **c,** Tumor cells (WR and non-WR), non-malignant astrocytes, oligodendrocytes and monocytes in multi-region sample SL040 as detected by EEL-FISH (full outline), corresponding MES-scores (dashed outline) and t-SNE representations of matched scRNA-seq data (circles). Low-fluorescence zone: sample SL040C; periphery: samples SL040F (top) and SL040D (bottom); high-fluorescence zone: samples SL040B (top) and SL040E (bottom).

We used droplet single-cell RNA sequencing (scRNA-seq) to collect 135,482 single-cell transcriptomes (after quality control (QC), Methods). We classified cells as malignant or non-malignant by inferred genomic copy number variations (CNVs) using infercnvpy^30^ (Methods, **Extended Data Fig. 1c**). We used established marker genes to identify astrocytes, B/plasma cells, endothelial cells, fibroblasts, macrophages, microglia, monocytes, oligodendrocytes, OPCs, pericytes, T/NK cells, tumor-associated macrophages (monocyte-derived, moTAMs), tumor-associated microglia (mgTAMs), tumor cells, vascular smooth muscle cells (Methods, **Fig. la, Extended Data Fig. 1d)**. We validated our myeloid cell annotation with a label transfer from an external dataset^31^. Tumor cells were highly frequent in the necrotic core and high-fluorescence zone (43 – 65% tumor cells). These two zones will henceforth be collectively referred to as the tumor bulk. Tumor cell frequency remained high in the low-fluorescence zone (25%, 65%), dropping sharply in the periphery, albeit with high variability across anatomical directions (0.2 – 10%; **Fig. la, Extended Data Fig. 1d**). The high frequency of tumor cells in the low-fluorescence zone, just outside the high-fluorescence zone (contrast-enhancing zone) is in line with previous findings^26^. Myeloid cells were also organized with high regional specificity: moTAMs in the necrotic core, a mix of moTAMs and mgTAMs in the high-fluorescence zone, microglia in the low-fluorescence zone, periphery and cortex (**Fig. la, Extended Data Fig. 1d**). The relative abundance of both healthy brain cell types (microglia and astrocytes) and tumor cells in the low-fluorescence zone likely reflects that tumor cell infiltration precedes shifts in other cell populations here. This, together with the corresponding high cell cycle scores (**Fig. 1a)**, implicates the low-fluorescence zone as a niche resembling ‘early-stage’ tumor growth, before bulk formation, substantiating the rationale behind ongoing trials of supramarginal resection of GBM^32,33^.

We found a strong regional specificity of the MES-like gene signature^5^ along the core-to-periphery axis, with highest expression levels in the tumor core (**Fig. 1a**, **Extended Data Fig. 1d)** in concordance with previous reports^34^. Outside the bulk, even in the low-fluorescence zone where tumor cell frequency remained high, MES-like cells were nearly absent (**Extended Data Fig. 1d**). Moreover, the expression of MES-like gene signature was not confined to malignant cells: astrocytes, oligodendrocytes and TAMs, among others, all exhibited varying expression levels of MES-like signature, universally increasing from periphery to core (**Fig. 1a,b**, **Extended Data Fig. 1e,f**). Nearly every gene of the MES-like signature was expressed and regulated in the same pattern in all cell types, and only a small set of genes (e.g., *GJA1* in astrocytes or *CLIC4* in oligodendrocytes) showed cell type-specific regulation (**Fig. 1b**). The most conspicuous exception was monocytes, where MES genes were generally not highly expressed or did not show regulation from core to periphery (**Fig. 1a**,**b****, Extended Data Fig. 1f**). We conclude that MES-like gene expression is a shared feature of many cell types and is not an intrinsic tumor cell state, but a spatially determined tissue state confined to the tumor bulk. We confirmed these findings on a second GBM case (123,236 cells after QC; **Extended Data Fig. 2a,c,d)** and on an external dataset^27^(3589 cells after QC; **Extended Data Fig. 2e,f**).

### Wound response as a generic feature of glioblastoma

The discovery that MES-like cell states are strongly regionalized and not tumor-specific prompted us to reconsider the interpretation of this gene expression module. The MES-like cell state in GBM was originally described based on bulk microarray gene expression profiles^1,8^. However, a gene ontology reanalysis of that data revealed no terms related to mesenchymal cells or processes, but rather to neural development and to wound healing, blood coagulation and apoptosis (**Supplementary Table 1**). Subsequently, Neftel *et al.* used single-cell transcriptomics to define two MES-like GBM cell states (MES1 and MES2)^5^. Gene ontology analyses of these gene sets returned no terms related to mesenchymal states or processes; instead, common significant terms were related to complement activation, plasma membrane repair, wound healing and coagulation (**Supplementary Table 1**). Namely, many MES1 genes are functionally involved in complement activation, plasma membrane repair, plasminogen activation and coagulation, all of which are related to hemostasis, i.e. the extravascular clotting of extravasated blood or plasma. For MES2, we found terms related mainly to hypoxia and none related to mesenchymal states or processes. A considerable fraction of MES2 genes are known direct targets of HIF1α, the master regulator of hypoxic responses (16 of 50 genes). Furthermore, all four Neftel *et al.* tumor cell states maintained the expression of glial master transcription factor *SOX2*, whereas functional mesenchymal marker genes (collagens, epithelial-to-mesenchymal, i.e. EMT, transcription factors) were either not expressed or expressed at low levels and not specific to MES-like cells (**Extended Data Fig. 3a**). Thus, these previously described MES-like gene sets seem neither mesenchymal nor specific to tumor cells. Instead, a detailed re-examination points to hypoxia, inflammation, injury, wound healing, blood clotting, and similar processes (**Supplementary Table 1**); that is, to a reactive wound-healing tissue state rather than a mesenchymal differentiation process. We therefore propose that the previously named mesenchymal-like cell states in GBM more likely represent fundamentally glial (*SOX2^+^*) tumor cells but with strong activation of hypoxia and wound healing responses analogous to those found in normal astrocytes during brain wound repair^35^.

To substantiate the hypothesis of MES gene involvement in wound healing responses, we examined their temporal response in three tumor-free injury settings. First, in a well-powered time course study of mouse skin wound healing^36^ (**Extended Data Fig. 3b**), all but one MES-core^37^ gene was robustly and significantly (FDR < 0.05, fold-change > 1.2) induced in the acute phase of wound response; *S100a16* was instead repressed in the late phase. Second, 23 of 28 MES-core genes were significantly (FDR < 0.05) induced in astrocytes in a mouse spinal cord injury model^38^ (**Extended Data Fig. 3c**). Finally, in human brain tissue resected 8 – 180 hours after traumatic brain injury^39^, 17 MES-like^5^ genes were significantly induced in astrocytes (four were down-regulated; n=12 patients and 5 controls, P < 10^−7^ and absolute log_2_ fold change > 0.25 as reported in the original publication; **Extended Data Fig. 3d**).

Curiously, we noticed that cells in the tumor bulk expressed far more mRNA in total, as compared with cells from the periphery: this was significant in tumor cells, TAMs, astrocytes, vascular cells, but not in oligodendrocytes (**Extended Data Fig 2b**). The mean number of reads per cell in bulk and periphery were similar across samples, but samples from bulk were generally less saturated despite a higher number of detected genes and unique molecular identifiers (UMIs) per cell. To discover which genes contributed most to this increase in overall mRNA abundance, we performed differential gene expression analysis between tumor bulk and periphery (**Extended Data Fig 1f, 2d,f**). Nearly all MES-like genes were upregulated in the tumor bulk across cell types, even when normalizing for total mRNA count. Furthermore, of the top upregulated genes in the bulk, the majority were MES-like genes and many of the remaining genes were closely related. Thus, we conclude that gene expression in the tumor bulk is dominated by a strongly induced transcriptional program, across tumor cells and most non-tumor cell types, that comprises mainly MES-like genes and closely related genes representing a wound healing response. We will therefore henceforth refer to them as ‘wound response’ (WR) cell states. We computed a consensus WR gene signature across cell types and samples (**Supplementary Table 2**). Reproducibility per cell type between samples was remarkably high with almost 40-50% overlap in the top ranking 500 genes, with the mismatched genes usually appearing in the following 500 genes (top 1000 genes). In addition to previously defined MES-like gene signatures, we found 19 transcription factors associated with the WR state (top 500 genes), including cAMP response-element binding protein (CREB) family genes *ATF5* and *CREB3*, C/EBP family *DDIT3* (the only TF in the previous MES-like signature), immediate-early genes *EGR2* and *FOSL2*, and *NFIL3* which controls expression of the inflammatory cytokine IL3. These factors provide a gene regulatory basis for activation of the WR phenotype in diverse cell types. Based on the enriched gene sets and our gene ontology analyses, it is likely that additional related processes such as inflammation, apoptosis, and metabolic rewiring also characterize these cellular states.

### Developmental and wound response states account for tumor cell diversity

To further investigate the nature of WR states, we expanded our cohort with standard samples of the bulk and mainly the contrast-enhancing tumor region, which largely correspond to the high-fluorescence zone and where WR states should be most abundant (totaling 25 GBMs and 2 oligodendrogliomas; **Supplementary Table 3**). We trained a graph attention neural network (GAT) to identify common molecular patterns across all samples by gene expression and spatial proximity (**Extended Data Fig. 5a**). The GAT output was clustered into molecular states (m-states) that distinguished microscale regions throughout the tissue, corresponding to cell transcriptional states or cell types. Cell segmentation by m-state yielded single cells (m-cells). In total, 42 shared m-states in 6,463,910 segmented m-cells (after QC; Methods) were identified from 11,081,393 imaged nuclei (**Extended Data Fig. 5a**). For initial annotation, we used Tangram^40^ to transfer cluster labels from a GBM scRNA-seq meta-analysis^41^. Most m-states mapped to either vascular, immune (validated by flow cytometry; **Extended Data Fig. 4a,b**) cell types or the four canonical tumor cell states (**Extended Data Fig. 5b**). To better distinguish non-malignant astrocytes from tumor AC-like cells, we trained a classifier on the matched scRNA-seq and EEL-FISH data from multi-region sample SL040. After confirming that the classifier could discriminate between astrocytes and AC-like m-cells in unseen sample with multi-regional representation (**Extended Data Fig. 4c**), we applied it to the remainder of the dataset. Most cell types showed strong core-to-periphery dichotomy, in agreement with our scRNA-seq data: oligodendrocytes and astrocytes were found primarily in the periphery, while tumor cells, despite infiltrating beyond the tumor bulk, were found in WR state predominantly within the high-fluorescence zone and only in small, sparse patches outside of it, matching the spatial distribution of the MES signature ( **Fig. 1c**). Thus, despite reports of MES-like state association with recurrence^8,9^, such cells were almost completely absent from the peripheral tumor regions that remain after resection.

Since the molecular diversity captured in our dataset exceeded the conventional nomenclature, we expanded our m-state classification using a recently published scRNA-seq dataset covering the first trimester of human brain development^42^ (**Extended Data Figure 5b**). We re-classified a total of 3,860,755 tumor m-cells by similarity to developmental cell types using Tangram, introducing glioblast (GBL)-like, neural intermediate progenitor cell (nIPC)-like, preOPC-like, and radial glia (RG)-like cell state labels. GBL-like and preOPC-like m-states were marked by the expression of *EDNRB* and *EGFR*^43,44^ (**Extended Data Fig. 6a**). In addition to the high *EGFR* levels, preOPC-like m-state also showed transcriptional resemblance to outer radial glia (*PTPRZ1*, *ITM2C*, *FABP7*, *EDNRB*), late glioblasts^42^ or the newly-described tripotential intermediate progenitor cells^45^, thus representing a less differentiated state than the initially assigned AC-like label (**Extended Data Fig. 5b**: II, **6a**). The nIPC-like m-states were defined by expression of *SOX2*, *MDM2, CNTN1* and/or *TCF7* (**Extended Data Fig. 6a**). Two m-states marked by expression of *SOX2*, *NES* and *VIM* mapped to radial glia and were accordingly named RG-like and RG-nIPC-like. The more ambivalent RG-nIPC-like classification stemmed from co-expression of *CDK4* and *GLI1*: the former has been linked to NPC-like cells in GBM^5^ and the latter is normally expressed in radial glia of the neocortex^46^. These newly annotated cell states generally coexisted with the canonical glia-like states and an abundance of WR cells (**Extended Data Fig. 6a**). Cases where glia-like m-cells vastly outnumbered WR m-cells (<5% of WR m-cells) were few but notably included the two oligodendrogliomas and mainly multi-region samples from outside the bulk (**Extended Data Fig. 6b**).

We subclassified WR m-states primarily by *VEGFA* expression levels and secondarily by other markers: *MDM2*, *PDGFRA*, *MGP, TNC* and *TIMP1* (**Extended Data Fig. 6a**). Multiple genes implicated in hypoxic and wound responses^47^ were characteristic of WR m-states: *VEGFA*, *LGALS1*, *SERPINE1*, *IGFBP5*, *ANXA1*, *MGP*, *TIMP1*. In contrast, we did not observe expression of true mesenchyme-specific genes such as type I or III collagens, suggesting that WR m-states were not of true mesenchymal identity. In part, the diversity of WR m-states reflected that of glia-like m-states: WR OPC expressed *PDGFRA,* a marker of OPC-like states, whereas WR nIPC expressed *MDM2*, linking it to nIPC-like 3 m-state (**Extended Data Fig. 6a**). The expression of hypoxia-response genes and the absence of mesenchymal markers in WR m-states supports the reactive state hypothesis; the transcriptional similarities might furthermore inform on the underlying glia-like identities.

### Wound response state is distinct from a fibroblast-like identity

To compare the newly-proposed WR and the conventional MES-like states, we examined the expression of mesenchymal and fibroblast markers, EMT transcription factors and the glial transcription factor *SOX2* in the original GBM cell state classification by Neftel *et al*.^5^ (**Extended Data Fig. 3a**). All malignant cell states maintained *SOX2* expression, unlike true fibroblasts or mesenchymal stem cells surveyed as non-malignant references^48^. Mesenchymal, fibroblast and EMT markers, while detectable to varying degrees, were not specific to the MES-like cell state. In fact, a single sample, pediatric glioblastoma BT830, accounted for nearly all fibroblast-like gene expression (collagens *COL1A1*, *COL1A2*, *COL3A1*, collagen cross-linkers *DCN* and *LUM*, but not *COL2A1* which is specific to cartilage) in the dataset (**Extended Data Fig. 3a**). Furthermore, these cells expressed markers of neural crest stem cells (*SOX10* and *NGFR*) and developing meningeal fibroblasts (*FOXC1*). Thus, the mesenchymal characteristics in the conventional definition of the MES-like state seem specific only to a small subpopulation of cells.

We analogously identified three distinctly fibroblast (FB)-like m-states in our EEL-FISH data, characterized by such fibroblast markers as *DCN* and *COL3A1* (**Extended Data Fig. 6a**). These m-states were found predominantly, albeit not exclusively, in two samples of gliosarcoma, a well-known subtype of glioblastoma characterized by a glial and a sarcomatous component (SL016A and SL020; **Extended Data Fig. 7a**). The *DCN*^+^*TIMP1*^+^*COL3A1*^+^ FB-like 3 m-state (**Extended Data Fig. 6a**), the most fibroblast-differentiated in our classification, was found abundantly in both gliosarcoma samples, coexisting with WR m-states (**Extended Data Fig. 7b**). Unlike the FB-like m-states, none of the WR states or other major gliosarcoma constituents (AC-like and RG-nIPC-like m-cells) were gliosarcoma-specific. In sample SL016A, *COL1A1*^+^ FB-like m-cells were positioned around endothelial (*RGCC*^+^) m-cells, as could be expected from fibroblast-like cells, and were in turn surrounded by *VEGFA*^+^ hypoxic WR m-states **(Extended Data Fig. 7b**). This vasculature-centered organization, however, was not universal: the AC-like-dominated sample SL020 showed far less of this perivascular organization, with FB-like m-cells being more spatially segregated. Such hubs of FB-like m-cells, marked by *TIMP1* expression, were found in both gliosarcoma samples (**Extended Data Fig. 7c**). Taken together, it appears that FB-like states have previously been subsumed under the conventional MES-like state, which, in turn, can be split into: (1) a reactive WR state not specific to tumor cells, and (2) a rare fibroblast-like tumor cell population enriched in gliosarcoma.

### Wound response is an inducible and reversible cell state

To test our hypothesis that WR is a modular and reactive state, we performed *in vitro* perturbations of glioblastoma organoids (GBOs) derived from four GBM patients. Given the expression of hypoxia and hemostasis-related genes in WR-states, we selected hypoxia and blood plasma as two candidate inducers. We subjected GBOs to either condition, or a combination of both, for 72 hours to test the inducibility, and reverted to regular media and normoxia for another 72 hours to assess the reversibility of WR state activation, performing Xenium spatial transcriptomics analysis at several timepoints (Methods; **Fig. 2a**). We integrated data from all four GBO lines using scVI^49^ and found a range of glia-like states corresponding to those observed in tumor samples, but only few non-malignant glial, immune or vascular cells (**Extended Data Fig. 8a**). We also found two hypoxic response states (+HYP1 and +HYP2) that resembled our previously described WR m-states by expression of *ANXA1/2*, *CD44*, *EPAS1, HILPDA* and *VEGFA*, among other markers. Cell states were reproducible across the different GBO lines even when not correcting for batch effects (**Extended Data Fig. 8b**).

**Figure 2.**
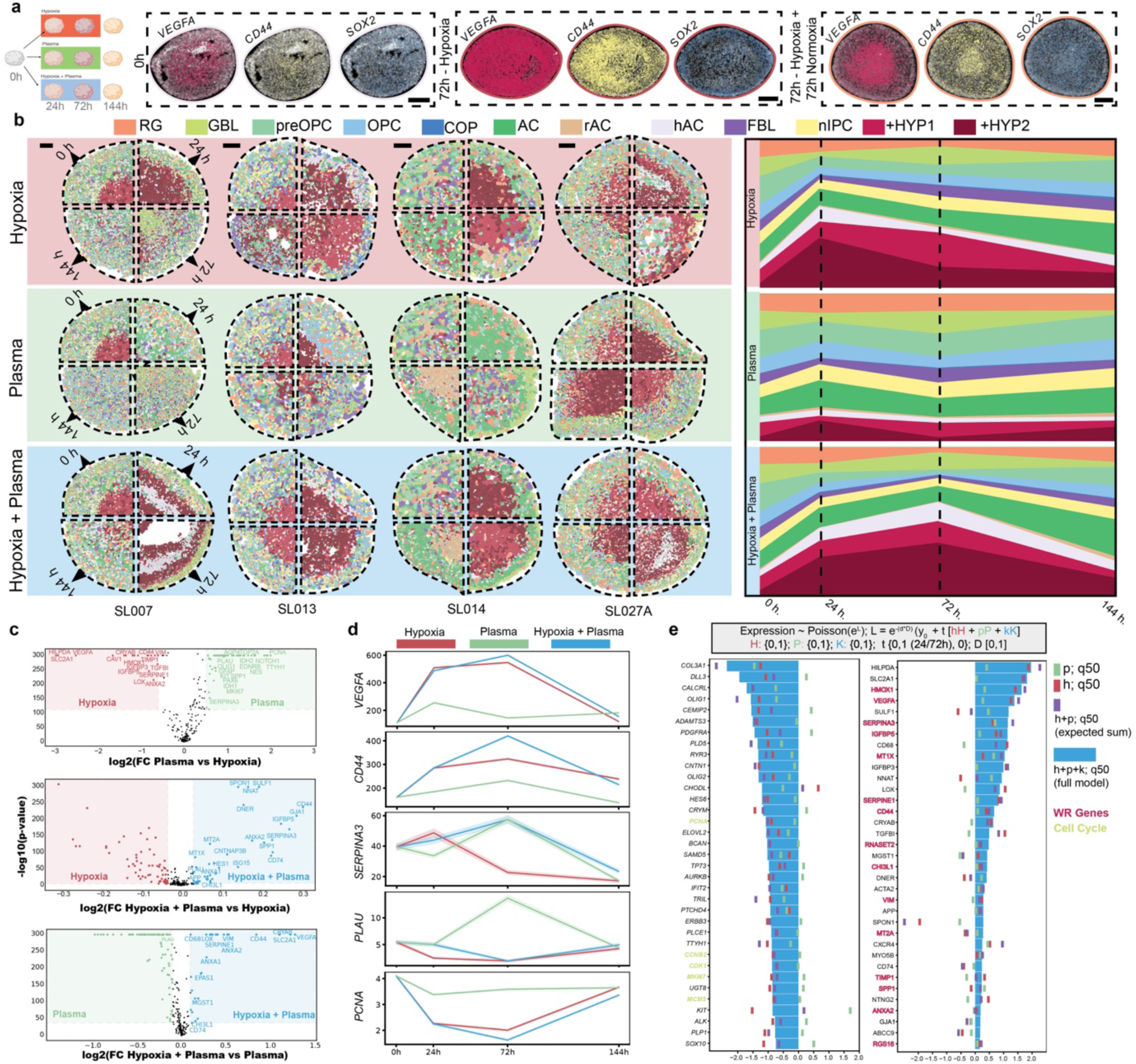
Transitional activation of WR program in patient-derived organoid lines. **a**, Left: schema of experiment. Right: expression of *SOX2*, *CD44* and *VEGFA* in organoid line SL027A over the course of experiment. Scale bar 200 µm. **b**, Left: spatial cell state distribution clocks of RG-, GBL-, preOPC-, OPC-, committed oligodendrocyte precursor (COP)-, AC-, reactive astrocyte (rAC)-, hypoxic astrocyte (hAC)-, FBL- and nIPC-like, +HYP1 and +HYP1 states at 0, 24, 72 and 144 hours when cultured under hypoxia, blood plasma or both. Scale bar (100 µm) corresponds to the top left quadrant (0 h timepoint), other quadrants are scaled to match the display size of the top left quadrant. Right: corresponding cell state proportions. **c**, Pairwise differential gene expression analysis between the different culturing conditions. **d**, Normalized expression of differentially expressed genes under different culturing conditions: *VEGFA*, *CD44, SERPINA3, PLAU* and *PCNA.* **e**, Top box: generalized linear model of gene expression according to a Poisson distribution with parameters: treatment timepoint (t), hypoxia (h), plasma (p), hypoxia and plasma (k), relative distance to organoid core (d). Left: genes with lowest effect size for k. Right: genes with highest effect size for combined treatment (k). Green line: estimated median from the posterior distribution for plasma effect size (p). Red line: estimated median from the posterior distribution for hypoxia effect size (h). Purple line: sum of hypoxia and plasma estimated medians. Blue bar: estimated median for the interaction effect between hypoxia and plasma (k). Cell cycle genes (green), wound response genes (red).

The glia-like states appeared spatially disorganized, whereas +HYP1 and +HYP2 states were localized in the center of the GBOs (**Fig. 2b**: left), as could be expected given an oxygen diffusion gradient along the surface-to-core axis. Both hypoxic programs were upregulated under hypoxia, reverting to starting levels upon return to normoxia (**Fig. 2b)**, whereas the glial marker *SOX2* maintained much more stable expression (**Fig 2a**). No pronounced changes in cell state proportion were associated with plasma exposure alone, and the effects of the combined hypoxia and plasma exposure paralleled those of hypoxia (**Fig. 2b**: right). Differential gene expression analysis of hypoxia and plasma treatments showed that hypoxia upregulated conventional MES-like genes (*ANXA2*, *CD44*, *HILPDA*, *VEGFA*, among others), whereas a more varied transcriptional signature was associated with plasma, including markers of proliferation (*PCNA*, *MKI67*, *TOP2A*), glial markers (*AQP4*, *GFAP*, *NOTCH1*, *NES*, *EDNRB*, *PAX6*), reactive gliosis marker *SERPINA3* and plasminogen activator *PLAU* (**Fig. 2c**: top). To account for inhibitory effects and synergistic interactions, we next compared the two individual culturing conditions to the combined one (**Fig. 2c**). Genes upregulated under combined exposure to plasma and hypoxia as compared to hypoxia alone included neurodevelopmental genes (*DNER, SPON1, SULF1, NNAT*), conventional MES-like markers (*ANXA1*, *CHI3L1*, *MT1X*, *MT2A*) and immune response-related genes (*SPP1*, *CD74*; **Fig 2c**: middle). Several of these (*ANXA1/2*, *CD44*, *CHI3L1*) were also upregulated in the combination treatment as compared to plasma treatment (**Fig 2c**: bottom), indicating a synergistic effect of hypoxia and plasma. Indeed, while for some genes the combined effect matched that of hypoxia (*VEGFA*, *PLAU, PCNA*) or plasma (*SERPINA3*) alone, others (*CD44*) responded to each condition differently (**Fig. 2d**).

To estimate the respective effect size of every factor on gene expression, we fit a Poisson regression model (**Fig. 2e**: top, **Extended Data Fig. 8c**). The combined hypoxia and plasma treatment upregulated multiple WR genes, and more strongly so than either condition alone (**Fig. 2e**: right). Conversely, cell cycle markers were among the most downregulated under combined treatment, although this effect was largely attributable to hypoxia alone (**Fig. 2e**: left). Curiously, in many instances, the true combined effect of hypoxia and plasma did not match the expected sum of effect sizes of each individual factor (*HILPDA*, *TGFBI*) or even opposed the expected sum effect (*SPON1, CD74, MGST1, MT2A, GJA1*, *ABCC9*). Taken together, these data show that a wound-like environment of hypoxia and blood plasma leads to a transient induction of WR signatures in tumor cells, supporting the reactive and modular nature of WR states in GBM.

### Vasculature is an organizing hub for wound response states

We next examined whether this reactive model of WR states could likewise help explain their organization in tumor context. We noticed that many tumors in our EEL-FISH dataset displayed a sharp structural dichotomy between WR regions and more peripheral areas containing non-WR tumor cells, astrocytes and oligodendrocytes (**Fig. 3a**). This distinction likely reflects the transition from the high- to the low-fluorescence zone described previously in the scRNA-seq data. Within the WR-rich zones, WR m-states were generally nested around blood vessels: an *MGP*^+^ WR m-state was associated with endothelial and mural cells (hence annotated as WR Endothelial Associated), *VEGFA*-high WR m-states (WR OPC, WR nIPC and WR HYP1/2) were located distally in respect to vasculature, and WR Periphery 1/2/3 m-states spread out from vessels to periphery (**Fig. 3a**: Ia,b, **Extended Data Fig. 7d**).

**Figure 3.**
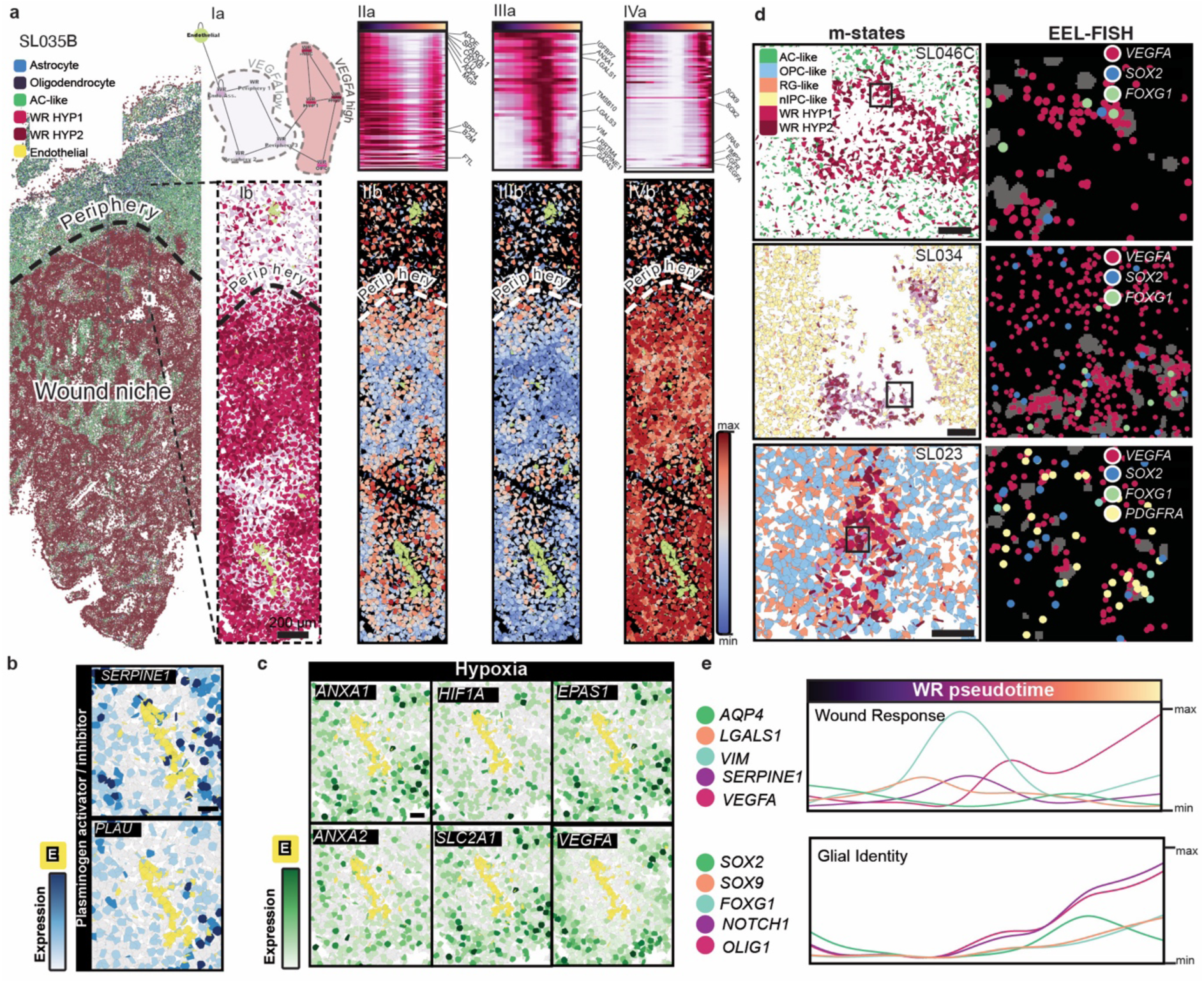
Activation of WR program in GBM. **a**, Left: spatial distribution of astrocytes, oligodendrocytes, AC-like, WR HYP1/2 and endothelial m-states in sample SL035B. Ia: Graph representation of spatial relationships between WR states and endothelial cells. Ib: Spatial distribution of WR and endothelial m-cells across peripheral, wound and perivascular niches in sample SL035B. IIa: peripheral/perivascular WR gene module expression along WR pseudotime; IIb: spatial distribution of the peripheral/perivascular WR gene module. IIIa: intermediate WR gene module expression along WR pseudotime; IIIb: spatial distribution of the intermediate WR gene module. IVa: wound niche WR gene module expression along WR pseudotime; IVb: spatial distribution of the wound niche WR gene module. **b**, Expression of *SERPINE1* and *PLAU* in the perivascular niche. Endothelial cells marked in yellow. **c**, Expression of *ANXA1/2*, *HIF1A*, *EPAS1*, *SLC2A1* and *VEGFA* in the perivascular niche. Endothelial cells marked in yellow. **d**, Spatial distribution of AC-like, OPC-like, RG-like, nIPC-like and WR HYP1/2 m-states in samples SL046C, SL034 and SL023. EEL-FISH co-expression for *VEGFA*, *SOX2*, *FOXG1* and *PDGFRA* in WR cells. **e**, WR pseudotime normalized expression of wound response (*AQP4*, *LGALS1*, *VIM*, *SERPINE1* and *VEGFA*) and glial (*SOX2*, *SOX9*, *FOXG1*, *NOTCH1* and *OLIG1*) genes.

We used scFates^50^ to fit a pseudotime trajectory along these m-states and observed a progression reflective of the spatial organization (**Extended Data Fig. 7d**). Namely, the wound response trajectory ranged from the less hypoxic WR m-states of the peripheral tumor regions (**Fig. 3a**: I, **Extended Data Fig. 7d**: SL002) to m-states with the highest *VEGFA* expression (WR HYP1/2), located farthest away from vasculature (**Fig. 3a**: I, **Extended Data Fig. 7d**: SL035B). We organized this WR state gradient into three gene modules that respectively defined the peripheral/perivascular (**Fig. 3a**: II), intermediate (**Fig. 3a**: III) and hypoxic states (**Fig. 3a**: IV). Notable hypoxia response genes (*ANXA1*/*2*, *HIF1A*, *SLC2A1*, *EPAS1*, *VEGFA*), signaling molecules *(NOTCH1, TGFBI, IGFBP5)*, and hemostasis regulators (*SERPINE1*, *PLAU*) followed a vasculature-centered gradient (**Fig. 3b,c**, **Extended Data Fig. 7e**), analogously to the surface-to-core axis in our GBO models. The structured transcriptional organization around vessels suggests that akin to *in vitro* induction in GBOs, WR states could be governed by tumor hypoxia gradients. In line with the reactive state hypothesis, we also found that WR m-cells maintained the expression of key glial markers (*SOX2*, *FOXG1* or *PDGFRA*) across tumors of distinct characteristic glia-like cell compositions (**Fig. 3d**). Expression of some glial factors (*SOX2*, *SOX9*, *FOXG1, OLIG1*, *NOTCH1*) even increased along the WR pseudotime axis (**Fig. 3e**). Thus, we conclude that WR activation does not negate glia-like identity of tumor cells and that the level of activation depends primarily on local hypoxia gradients, in line with recent reports identifying hypoxia as an organizing factor in GBM^51^.

### Hypoxia links immune infiltration to wound response microenvironment

We next asked what other cellular components contributed to the wound response microenvironment. Firstly, we found a particularly strong and widespread spatial association between hypoxic tumor WR m-cells (WR HYP), monocytes (Mono1 and Mono2) and hypoxic moTAMs (moTAM HYP, *SPP1^+^MT1H^+^*; **Fig. 4a**: I). Conversely, non-hypoxic moTAMs and mgTAMs were associated with peripheral WR and non-WR tumor m-cells, alongside non-malignant astrocytes and oligodendrocytes (**Fig. 4a**: I).

**Figure 4.**
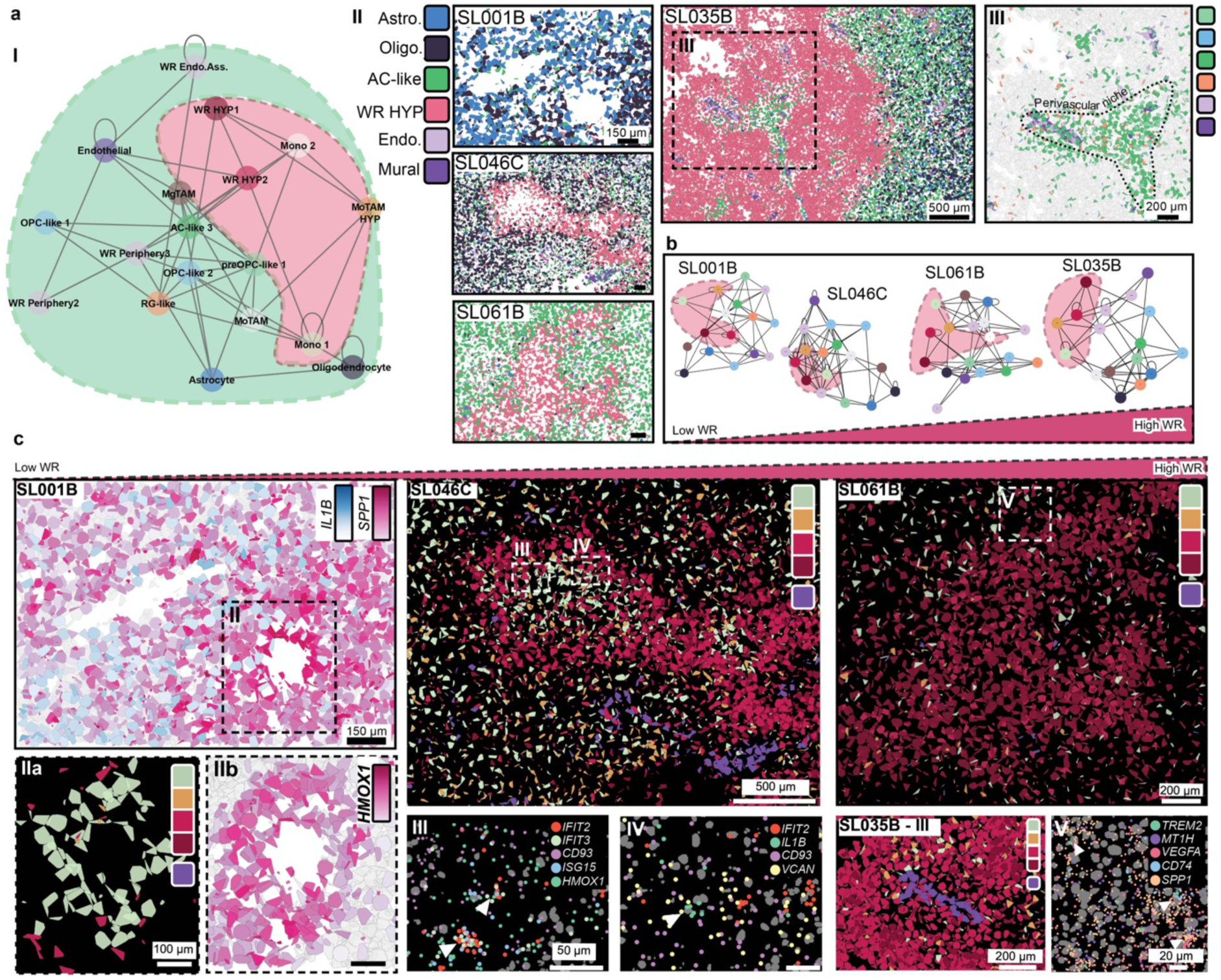
Immune infiltration of the WR niche in GBM. **a**, I: Spatial connectivity graph of GBM m-states across all samples. Edges indicate significant connections. Immune-WR niche highlighted in red. II: Spatial distribution of astrocyte, oligodendrocyte, AC-like, WR HYP, endothelial and mural m-states in samples SL001B, SL046C, SL061B and SL035B. III: spatial distribution of m-states in a perivascular niche in sample SL035B. Color legend matches panel I. **b**, Spatial connectivity graph of m-states in samples SL001B, SL046C, SL061B and SL035B. **c**, Left: IL1B and SPP1 expression in sample SL001B (top). IIa: Spatial distribution of monocyte, moTAM HYP, WR HYP, endothelial and mural m-states in sample SL001B. IIb: HMOX1 expression in monocyte infiltrating region of sample SL001B. Middle: spatial distribution of monocyte, moTAM HYP, WR HYP, endothelial and mural m-states in sample SL046C (top). III and IV: IFIT2/3, IL1B and ISG15 expression in hypoxic (HMOX1) and monocyte infiltrating (CD93/VCAN) regions of sample SL046C. Right: spatial distribution of monocyte, moTAM HYP, WR HYP, endothelial and mural m-states in samples SL061B and SL035B. V: anti-inflammatory (TREM2) and hypoxia signatures (SPP1, MT1H, VEGFA) in moTAM HYP infiltrating (CD74+) region of sample SL061B. M-state color legends match panel **a**(I).

To investigate how this immune-WR niche could be established, we selected four samples ranging from low to high WR content, thereby approximating a time series of the niche construction (**Fig. 4a**: II). The association between infiltrating monocytes (Mono1), moTAM HYP and tumor WR m-cells was maintained regardless of WR level (**Fig. 4b**). Notably, even in the most WR-rich samples, perivascular niches were not overridden by WR (**Fig. 4a**: III). Conversely, even in extremely low-WR tumors, we found areas of high expression of both the pro-inflammatory *IL1B* and the hypoxic *SPP1* (**Fig. 4c**: SL001B). These areas were infiltrated by monocytes (**Fig. 4c**: IIa), particularly in the more hypoxic pockets, as marked by high *HMOX1* expression (**Fig. 4c**: IIb), possibly reflecting an early stage of the immune-WR niche progression. Similar, but more advanced patterns were observed in tumors with higher WR content, where monocytes were mixed with moTAMs and surrounded by WR HYP m-cells (**Fig. 4c**: SL046C). Within the hypoxic pockets of monocytes, we located m-cells expressing interferon-related genes *IFIT2, IFIT3, ISG15* and *IL1B* (**Fig. 4c**: III,IV), similar to the previously reported signature of CD16^-^ monocytes^52^. In samples with high WR content, WR niches were far more scarcely infiltrated by monocytes, with similar levels of moTAM HYP instead (**Fig. 4c**: SL061B and SL035B). We believe that these data might indeed capture the gradual progression of WR niches, beginning with monocyte recruitment to hypoxic areas (*HMOX1^+^*) and a concurrent switch towards WR states in tumor cells, whereupon monocytes remain sequestered among WR tumor cells, differentiating into immunosuppressive macrophages (*CD74^+^CD163*^+^*TREM2*^+^) with a hypoxic signature (**Fig. 4c**: V). Such a sequence of events would be in line with the reports showing that hypoxic niches attract and sequester TAMs, ultimately leading to pseudopalisade formation^53^.

### Wound response-independent spatial organization is conserved across patients

We lastly searched for universal spatial GBM characteristics conserved between patients. Matching mutational profiles often corresponded to spatial profile similarity but the most striking division was found between WR-low and WR-high samples (**Fig. 5a**). The stratification by WR activation was observed even within a sample originating from the same patient (e.g., SL046B/C) suggesting that local WR effects, rather than mutational predispositions, dominate tumor niche architecture. We reevaluated spatial associations by excluding WR m-states to eliminate potential masking effects (**Fig. 5b**). AC-like 3 (*CRYAB*^+^) m-state was recurrently found near endothelial and mural cells (**Fig. 5b,c**, **Extended Data Fig. 9**), similarly to the well-documented colocalization of healthy mature astrocytes with vasculature^54^. We observed this association across tumors of varied WR proportions and mutation profiles (**Fig. 5c**: SL035B, SL034 and SL046B/C). Similarly, we observed a consistent association between preOPC-like 1 and OPC-like 2 m-cells (**Fig. 5b,c**). The spatial association between the supposed precursor-descendant state pair supports our classification of this *EGFR*^+^ m-state as preOPC-like. Taken together, these data suggest that the basic GBM organization, upon which WR activation is superimposed, retains features characteristic of its tissue of origin, the brain.

**Figure 5.**
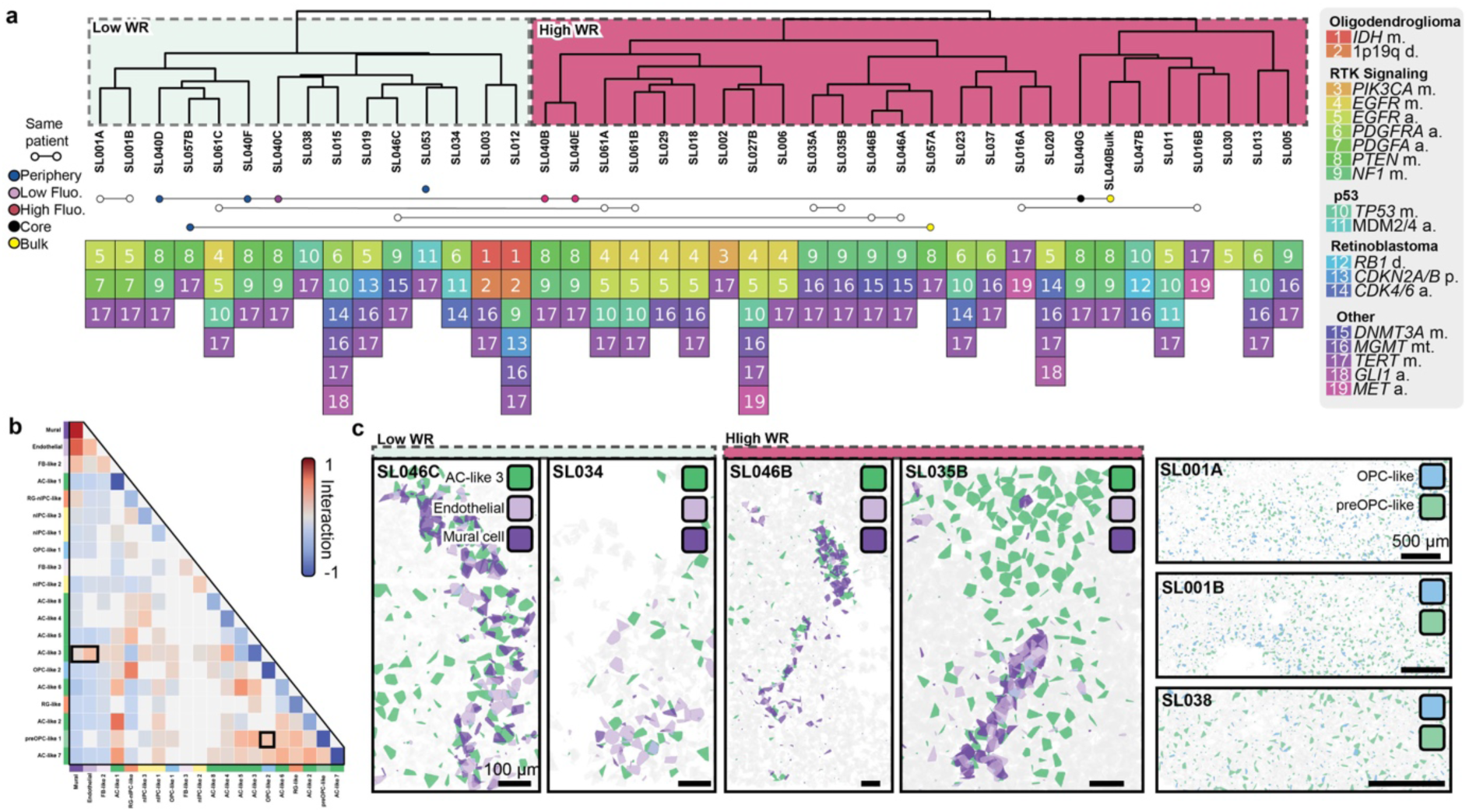
Spatial organization of cell states across a large GBM cohort. **a**, Dendrogram of m-state spatial organization similarity across samples with corresponding mutational profiles. Connected dots mark samples originating from one patient, color-coded by sampling location. Each square indicates mutations identified in the pathology report (Supplementary Table 3): mutated (m.), amplified (a.), deleted (d.), positive signal (p.), methylated (mt.). **b**, Neighborhood enrichment scores for non-WR tumor, endothelial and mural m-states in all samples. **c**, Spatial distribution of AC-like 3, endothelial and mural m-samples SL034, SL035, SL046B/C. Spatial distribution of OPC-like 2 and preOPC-like 1 m-states in samples SL001A/B and SL038. Gray polygons represent all other m-states.

## Discussion

Our understanding of GBM has been largely built on data acquired from the contrast-enhancing zone of the tumor bulk, as determined by the current clinical practices of maximal safe resection. By studying rare cases where tissue from tumor periphery was also accessible, we showed that the molecular architecture changes drastically along the core-to-periphery axis. The tumor bulk was dominated by a hypoxia and wound response program shared across malignant and non-malignant cell types and an abundance of TAMs. In contrast, the low-fluorescence zone just outside the tumor bulk showed very little wound response and contained normal glia and lack of TAMs despite considerable infiltration by proliferative tumor cells. We believe that the emergence of WR states in the tumor bulk may reflect a fundamental self-perpetuating mechanism in GBM (**Extended Data Fig. 10**): a growing tumor causes tissue damage, vascular dysfunction and oxygen depletion, inducing hypoxia and clotting that activate WR states, which, through secreted factors such as *VEGFA*, further increase vascular leakage and thus reinforce WR activation in a cycle of futile wound response^55^. Monocytes recruited to this unresolvable wound niche would ultimately differentiate towards WR-signature expressing moTAMs and further contribute to the wound-like environment. Metabolic factors, like high oxygen consumption by myeloid cells^56^ could further reinforce the vicious cycle. While intrinsic factors, namely *NF1* alterations, have been linked to the MES-like state^1,5^, our multi-region sample SL040 lacked MES-like signature expression outside of the tumor bulk despite carrying a clinically confirmed *NF1*-mutation (**Supplementary Table 3**), showing that *NF1* alterations were insufficient to intrinsically induce MES-like states outside of their requisite spatial context. This re-interpretation of MES-like (*i.e.*, WR) phenotypes not as tumor-intrinsic but rather as tissue-wide reactive states driven by rapid tumor growth would imply that these cell states are a consequence of tumor aggressiveness, not its cause.

Importantly, such chronic and hypoxic wound response mechanisms might not be GBM-specific: multiple studies in solid tumors have shown that hypoxic gene signatures are prognostic markers of an aggressive disease with a short survival^57–60^. Five of the top 15 genes in the shortlist from various solid tumors by Buffa *et al*.^58^ are part of the MES-like signature (S*LC2A1, LDHA, P4HA1, ADM, NDRG1*), and another six are present in our WR signature (Supplementary table 2). Of the 15 genes shortlisted in pancreatic cancer by Khouzam *et al*.^60^, nine are part of the MES-signature (*NDRG1, SLC2A1, P4HA1, LDHA, PGK1, ANGPTL, BNIP3, ADM, CA9*).

In summary, our findings show that the WR signature is cell-extrinsic, confined to the bulk of the tumor and shared across tumor and non-tumor cell types. Given our findings of WR activation as a strong central and universal feature of GBM, other recent reports highlighting the role of injury-like processes in GBM progression^61^ and the suspected relevance of WR in other solid tumors, the reconceptualization of MES-like cell states as WR is not a simple refinement of nomenclature but a shift in perspective regarding the interplay between tumor cell type, state and tissue organization. Recognizing WR as a modular tissue state should spark renewed interest in the truly tumor-intrinsic malignant cell types and the unique microenvironment outside the tumor bulk, the probable source of most relapses in GBM.

## METHODS

### Patient samples

Human glioblastoma samples were collected with informed consent from the patients and with ethical approval from the Swedish Ethical Review Authority (2020-03505). The use of samples was approved by the Swedish Ethical Review Authority (2020-02096). Information on clinical metadata and neuropathology data can be found in Supplementary Table 3.

### Human tissue collection

GBM samples were kept fresh in Hibernate-E medium (Gibco) on ice during transportation from neurosurgery at the Karolinska University Hospital. Upon arrival, samples were either directly processed for scRNA-seq or embedded in Tissue-Tek Optimal Cutting Temperature (O.C.T.; Sakura) compound and snap-frozen in 2-methylbutane (Sigma Aldrich) and dry ice slurry. Frozen samples were stored at −80°C until further use.

### Multi-region sampling rationale

Neuronavigation, using preoperative MRI, was used according to standard of care. This is a central tool used on a regular basis in surgery of GBM. However, brain shift is a notorious and frequent problem and increasingly so during the continuous surgical removal of the tumor^62^. This compromises the accuracy of biopsies, especially when sampling from multiple locations. The fluorescence from 5-ALA, on the other hand, corresponds very well with the contrast-enhancing zone and extends beyond its outer margin, enabling an intraoperative delineation and guidance for the neurosurgeon^63^. Sampling based on 5-ALA fluorescence eliminates brain shift bias and we therefore decided to define the location of the biopsies based on the fluorescence rather than neuronavigation alone. Systematic sampling was done at multiple distances from the center of the tumor core in four different directions (anterior, posterior, superior, inferior) using a biopsy rongeur yielding approximately 3×5 mm tissue biopsies from each location. Sampling was performed early in the resection procedure to minimize brain shift, and each sample individually analyzed under blue light to define fluorescence intensity.

### Tissue dissociation

First, the tissue was chopped into small pieces using a razor blade. The Worthington’s Papain Dissociation System (Worthington) was used for enzymatic digestion at 37°C for 15-40 min (depending on the region, cortical samples usually required longer time). Manual trituration of the suspension was performed with glass pipettes. The suspension was subsequently filtered through a 70-µm filter followed by a 30-µm filter (Miltenyi, CellTrics). After centrifugation (200 g for 5 min at 4-8°C) the pellet was resuspended in EBSS (Worthington). Gradient centrifugation was performed using an albumin inhibitor solution (Worthington), followed by lysis of erythrocytes with RBC lysis buffer (Invitrogen) according to the manufacturer’s instructions. After final resuspension in EBSS visual inspection was done on a hemocytometer to estimate viability of the suspension (1 part Trypan blue 0.4%, Gibco, and 1 part cell suspension) and confirm minimal presence of multiplets, debris and to estimate the concentration of cells.

### Single-cell RNA sequencing

We processed two to three replicates per sample with the 10X Genomics V3.1 kits. If needed, dilution or resuspension was performed to reach a concentration corresponding to a target of about 10,000 cells per replicate when loading on the 10X Genomics chromium chips. Shallow sequencing (about 1000 reads per cell, Illumina NextSeq platform) was performed for quality control and estimate of cell concentrations followed by deep sequencing of about 100,000 reads per cell on the Illumina NovasSeq platform.

Sequencing runs were demultiplexed, filtered through an index-hopping filter, UMIs were counted and reads aligned to a human reference genome as previously described^42^.

### Single-cell RNA sequencing quality control

Broken 10X chromium emulsions were discarded. After library preparation, the cDNA yield and fragment distribution were evaluated (2100 Bioanalyzer Instrument, Agilent Technologies). Replicates with more than 25,000 cells after deep sequencing were not analyzed further. Droplet classification was based on a previously described approach^42^; thresholds were modified based on evaluating the droplet distribution (UMI vs unspliced read fraction) with the final criteria: minimum UMI count of 4000, minimum unspliced read fraction of 0.1, maximum doublet score of 0.4, maximum mitochondrial read fraction of 0.25, two-dimensional Gaussian maximum likelihood values m=30, k=1500.

### Single-cell RNA sequencing data analysis

Data were analyzed using Cytograph-Shoji (https://github.com/linnarsson-lab/cytograph-shoji), which implements a custom tensor database. All cells that passed our QC criteria were clustered as previously described^42^ but with 2000 most variable genes (instead of 1000) selected using support-vector regression on the CV-versus-mean relationship. Harmony^64^ was not required since only one 10X chemistry version was used.

### Single-cell annotation

Manual curation was performed based on multiple established cell type-specific markers. Cells with combined expression of genes specific to more than one cell type were suspected to be doublets and were filtered out. After this step, re-clustering and re-annotation was performed. For distinction within the myeloid populations the following genes were used. mgTAMs shared expression of microglia genes (*CX3CR1, P2RY12, P2RY13, TMEM119*) as well as moTAM genes (*F13A1, S100A8, S100A9*) which were absent in microglia. For mgTAMs *HAMP* was specific. Microglia was the only cell type expressing the dendritic cell markers *CLEC9A* and *FCER1A*. Genes present in both macrophages and moTAMs were *ITGA4, RNASE1, F13A1* but macrophages lacked TAM genes (*S100A8, S100A9, SPP1, VCAN)* and *EREG,* a gene specific for a subset of moTAMs.

### Inferring CNVs in single cells

We used Infercnvpy^30^, a Scanpy^65^ plugin, to infer CNVs from scRNA-seq data in order to identify tumor cells. Data were normalized and logarithmized (base *e*) in Scanpy. Window size was set to 500 genes. Identified tumor cells matched and validated our annotation of tumor cells based on gene expression. Cells from various non-malignant cell types were then used as reference categories to set the background CNV profile.

### Meta-module scoring

The calculation was performed according to the algorithm described by Neftel *et al.*^5^ The code was implemented in Python and first validated on the dataset from Neftel *et al.* to ensure the results matched the original publication. Every cell has four different scores, each representing the substates (including the MES-state score).

### Differential gene expression

Data were normalized and logarithmized in Scanpy^65^. The Scanpy function “rank_genes_group” was used to rank genes according to Z-score (Wilcoxon) and the corresponding significance-level (p-value). Genes were plotted according to its scores (ranking top 20 genes) and visualized in a volcano plot. Genes that are part of the MES-genes according to Neftel *et al*. were encircled in red.

### EEL-FISH probe design

The RNA-binding encoding probes were designed to previously reported specifications^12^ using Oligopy (https://github.com/linnarsson-lab/oligopy). Gene identities were encoded in 16-bit binary barcodes, with each barcode containing six positive bits to be read out in a single fluorescence channel, thus allowing up to 448-plex gene detection per fluorescence channel over 16 rounds of imaging. In total, we used three different encoding probe panels, each designed to be read out in different fluorescence channels for orthogonal detection. Two of these panels constituted the 888 genes used in further analysis; the third panel was used only as a back-up for a subset of the 888 main genes.

Encoding probes (produced as Oligo Pools, Twist Bioscience) were amplified in-house. Probe amplification was carried out separately for each of the three probe sets, following a previously published protocol^66^. A modification was made to the original protocol in the reverse transcription reaction: a 5′-amine-modified forward primer was used to improve encoding probe anchoring during formaldehyde fixation in the EEL-FISH protocol. The RNA input concentration for the reverse transcription step was limited to approximately 150 ng/μl.

Readout probes (IDT) contained 20-nucleotide-long sequences complementary to the detection sequences on the encoding probes and a disulfide-conjugated fluorophores (Cy3, Atto 590 or Alexa Fluor 647) on either 5’- or both 5’- and 3’-ends.

All probe and primer sequences are available in Supplementary Table 4.

### EEL-FISH

EEL-FISH was performed as previously described^12^, with a digestion buffer adapted for glioblastoma tissue. Namely, the digestion buffer contained 1% (wt/vol) sodium dodecyl sulfate (SDS, Sigma-Aldrich), 20 mM tris-HCl pH 7.4 (Thermo Fisher Scientific), 100 U/ml Superase RNase inhibitor (Thermo Fisher Scientific), and 0.8 U/ml proteinase K (NEB). The samples incubated with the modified digestion buffer four times for five minutes at 30°C, followed by four washes with pre-warmed 5% (wt/vol) SDS in 2× SSC (Sigma-Aldrich) for five minutes each at 30°C. We noticed high variability in glioblastoma tissue sensitivity to the digestion treatment and therefore we strongly recommend adjusting the proteinase K concentration and the number of digestion buffer washes for each sample individually.

Tissue sections adjacent to EEL-FISH section were used for H&E (Hematoxylin and Eosin stain kit; Vector laboratories; see https://protocols.io/view/hematoxylin-amp-eosin-staining-protocol-djrr4m56 for full protocol).

### EEL-FISH image acquisition

EEL-FISH image acquisition was performed on a Nikon Ti2 epifluorescence microscope equipped with a Nikon CFI Plan Apo Lambda ×60 oil immersion objective with a numerical aperture (NA) of 1.4, CFI Plan Apo Lambda ×40 objective with NA 0.95, CFI Plan Apo Lambda ×10 with NA 0.45, Nikon motorized stage, Nikon Perfect Focus system, Sona 4.2B-11 back-illuminated sCMOS camera with a pixel size of 11 μm (Andor). We used Lumencor Spectra light engine and matching filter cubes from Chroma. Namely, fiducial europium beads were imaged using 360/28 bandpass filter (400 mW light source; 30% power), ET667/30m emission filter, and T647lpxr dichroic mirror; Cy3 channel was imaged using 534/20 bandpass filter (400 mW light source, 100% power), ET575/40m emission filter, and T550lpxr dichroic mirror; Atto 590 channel was imaged using 586/20 bandpass filter (400 mW light source, 100% power), 89403X excitation filter, 89403m multiband emission filter, and 89403bs dichroic mirror; Alexa Fluor 647 channel was imaged using 631/28 bandpass filter (500 mW light source, 100% power), ET667/30m emission filter, and T647lpxr dichroic mirror, with all exposure times set to 100 ms.

Histology image acquisition was performed at Karolinska Institutet’s Biomedicum Imaging Core facility on a Zeiss Axio Scan.Z1 slide scanner.

### Flow cytometry

Samples were minced with a scalpel into ∼1-3 mm pieces and digested in RPMI-1640 medium (Cytiva) containing 0.25 mg/ml DNase I (Roche) and 0.25 mg/ml Collagenase (Sigma-Aldrich) at 37 °C for 25 min with stirring, strained through a 70-µm strainer and neutralized by adding RPMI-1640 with 10% heat inactivated FBS (HI FBS; Gibco).

Flow cytometry was performed as previously described^67^. Briefly, cells were stained for 20 min at RT with extracellular antibodies in a staining mix containing FACS buffer (1× dPBS (Sigma-Aldrich) with 2% HI FBS and 2 mM EDTA (Invitrogen)), 1:5 BD Horizon Brilliant Stain buffer (BD Biosciences), and 1:25 FcR blocking reagent (Miltenyi), followed by fixation using BD FixPerm buffer (BD Biosciences) for 30 min at RT, and staining with intracellular antibodies in BD Perm/Wash buffer (BD Biosciences) for 20 min at RT. Flow cytometry was performed on a FACSymphony A5 (BD Biosciences), fcs files generated using DIVA (BD Biosciences) and data analyzed in FlowJo v10.5.3 (BD Biosciences).

### Glioblastoma organoid culture

Organoids were grown as described in Jacob *et al.*^68^ from patient samples described above. Organoids were biobanked in liquid nitrogen at passage 0-1, revived and grown to passage 3-4. Organoids of homogenous size and shape were selected for experiment and grown in 4 ml medium per condition in 6-well plates with continuous shaking. For blood plasma condition, medium containing 10% human plasma (Merck) was prepared by decreasing the volume of DMEM/F12 and Neurobasal medium, while the concentrations of supplements were identical to the standard medium. For hypoxia condition, organoids were grown in moderate hypoxia (5% O_2_, 5% CO_2_) for the first 24 h and severe hypoxia (2% O_2_, 5% CO_2_) from timepoint 24 h to 72 h.

### Xenium sample preparation

Organoids were washed with 1× PBS (Thermo Fisher Scientific), fixed in 4% formaldehyde (Thermo Fisher Scientific) in 1× PBS for 10 minutes and washed again with 1× PBS three times. Before freezing, organoids were cryoprotected by allowing to sink in 15% and then 30% sucrose (Sigma Aldrich) solutions in 1× PBS. Xenium (v1 platform, Human Brain panel with custom add-on genes; 10x Genomics; Supplementary Table 4) experiments were performed by the In Situ Sequencing Infrastructure Unit, SciLifeLab Stockholm.

### Image processing

We used pysmFISH (https://github.com/linnarsson-lab/pysmFISH_auto) to automatically detect and decode the EEL-FISH signal. Briefly, after parsing, images were filtered, and large objects with a size over the 95^th^ quantile were removed. Then, spots were counted and registered between hybridizations using europium fiducial beads. The barcodes were then identified using a nearest-neighbor approach. To map the EEL-FISH signal (×60 objective) to nuclei images (×40 objective), we first registered the europium beads imaged with both objectives by using a point set alignment algorithm based on a nearest-neighbor search, and the resulting transformation (scaling, rotation and shift) was applied to the EEL-FISH signal. Nuclei were segmented using Cellpose^69^. To process the large amount of data generated by each EEL-FISH experiment, the analysis is parallelized using Dask^70^. EEL-FISH data were aligned to histology images based on nuclei masks using SimpleElastix (affine transformation with 15 resolutions and 2500 iterations; https://github.com/SuperElastix/SimpleElastix).

### M-state Graph Attention Network

Transcriptional States, m-states, were inferred using a cell-free segmentation approach using a graph attention neural network (GAT) as previously proposed by Partel and Whalby^71^. Only the 888 RNA species common to all EEL-FISH experiments were used for analysis. *GFAP* and *MOG* molecules were removed due to their extremely high counts. First, molecules were connected into an undirected graph if the distance between two molecules was below the 95^th^ quantile at which all molecules were connected to at least 3 neighboring molecules (∼5-10 μm). Nodes in the graph represent RNA molecules and every node has a feature vector which is a one hot encoding the identity of the RNA transcript, edges in the graph connect neighboring molecules. We generated a graph for each EEL-FISH sample and pruned all graphs by removing molecules connected to less than 5 neighbors. The GAT was designed using PyTorch (https://pytorch.org) and Deep Graph Library (https://www.dgl.ai/) with the following architecture: 2-hop neighbor aggregation using GAT2vCONV^72,73^ module from DGL and a final linear layer; and the following hyperparameters: a random sampling strategy with 20 and 10 neighbors for 1^st^ and 2^nd^ hop, respectively, 64 units for each hidden layer and 48 units for generated lower dimensional embedding. The GAT was trained using Adam as optimizer^74^ with a learning rate of 0.0001 and mini-batch sampling of 512 nodes per iteration. Model parameters are learned using an unsupervised method under the premise that neighboring RNA molecules in the should have similar embeddings, whilst distant molecules should have dissimilar embeddings. During training, embeddings are generated for *positive* nodes, true neighbors, and *negative* nodes, randomly sampled nodes, from each central node and the model’s objective function consists of the binary cross-entropy between *positive* and *negative* embedding for each sampled node.

We trained our generalized GBM-GAT for 10 epochs on 2-hop subgraphs generated from 500,000 random nodes, from 30% of our EEL-FISH sections, constituting a graph for training with a total of 33.6 million nodes and 359.7 million edges. The generated embedding for all central nodes was clustered into 80 different m-states using sklearn function *MiniBatchKMeans* and a sklearn *SGDClassifier* was trained using the generated embedding to predict cluster id from GAT embedding vectors. Next, we applied our generalized GBM-GAT to each individual EEL-FISH section graph and classified each RNA into the 80 spatial domains using the trained classifier. RNA molecules with a prediction probability below 50% to a spatial domain were discarded. Finally, we reduced m-states to 42 after merging m-states with gene expression correlation over 85%.

### M-cell segmentation

To facilitate further analysis, m-states were segmented using a combination of DBscan and QTclustering. First, sklearn function *DBscan* was applied to each individual m-state using 20 and 10 for the parameters_*eps* and *min_samples*, respectively. Next, we re-segmented DBscan outputs, if they had a maximum distance over 50 μm, using QTclustering algorithm with parameters *max_radius* and *min_cluster_size*, 20 and 10, respectively. We removed m-cells with less than 10 molecules. Finally, m-cells were kept if the distance to closest cell nuclei was below 50 μm.

### M-state annotation

Annotation of m-states was done by combining Tangram^40^ predictions and gene enrichment scores. First, we used Tangram to map all m-states gene expression from all samples to GBmap^41^ averaged expression of tumor states, immune cells, and oligodendrocytes. Gene expression from each m-state was normalized to 10,000 molecules per m-cell and Tangram was run using the following parameters: *mode=*’clusters’, *num_epochs*=1000, *density_prior*=’uniform’ and *lambda_g2*=1. The probability of each m-state for each reference cluster was averaged for all samples. The same procedure was repeated to map the m-states to human developmental scRNA-seq data^42^.

Gene enrichment was calculated for every m-state and sample using Cytograph’s enrichment score function *Enrichment*.

Multi-region sample SL040 was used to re-classify AC-like m-cells as either non-tumor astrocytes (periphery samples) or AC-like tumor astrocytes (high-fluorescence zone). A linear classifier (CellTypist^75^) was trained on samples from the periphery or high-fluorescence zone from sample SL040. The classifier was applied to the remaining EEL-FISH samples to reclassify non-tumor astrocytes m-states accordingly.

### scVI integration of organoid data

We utilized scVI^49^ to model gene expression profiles from xenium organoid data. We trained two models, with and without patient-line correction, to account and assess the biological and technical variability that arise from using 4 different patient GBM organoid lines, 3 different conditions (hypoxia, blood plasma, and both), and 4 different time points (0, 24, 48 and 144 hours). Leiden clustering was used to identify common molecular profiles across all conditions and patients using the batch-corrected latent space from scVI.

### Generalized linear model of Wound Response activation-deactivation in organoids

We implemented a generalized linear model (GLM) in PyMC^76^ to model the expression of each gene as a Poisson-distributed variable, using Markov chain Monte Carlo (MCMC) to infer the posterior distributions for the effect sizes of hypoxia, plasma, their interaction, and distance to core (eq1. and eq. 2); where β, d, h, p, and k are the intercept, effect size of the distance to core, hypoxia, blood plasma, and interaction effect for hypoxia and plasma, respectively. We used non informative normal priors with a zero-centered mean and standard deviation of 5. Time (t) was indicated as either 0 (for time 0 h and 144 h) or 1 for time 24 h or 48 h under hypoxia, plasma or both. Distance to core (D) was normalized between 0 and 1. Posterior distributions were inferred using PyMC NUTS sampler.

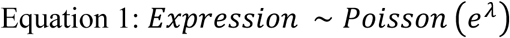

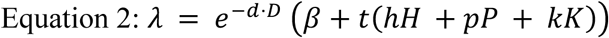

### Wound Response pseudotime

We used scFates^50^ to infer a pseudotime for the activation of wound response on the EEL-FISH data. We used the m-states identified as WR from all EEL-FISH samples, filtered cells with less than 5 molecules, normalized, log-transformed, scaled, and obtained the principal components using only highly variable features. The scFates function sc.tl.curve was used to build a graph using the following hyperparameters (*ndims_rep=2, epg_mu=.5*). The root was defined as the closest to highest *AQP4* and lowest *VEGFA* expression. Next, cells were projected into the pseudotime, and genes were clustered into different modules/trends.

### Neighborhood enrichment and graphs

Squidpy^77^ functions were used to calculate the enrichment scores for all possible m-state pairs. First, *spatial_neighbors* function was used to build a neighborhood graph connecting each m-cell to the 10 nearest m-cells below 100 μm distance and used Delaunay triangulation. Then, we used *neighborhood_enrichment* to obtain the spatial proximity scores for each pair based on permutation test.

We calculated a spatial similarity dendrogram based on each tumor sample enrichment score. First, enrichment score matrices were vectorized and rearranged so that each vector entry corresponded to the enrichment score for the same m-state pair in each sample. Next, each vector was scaled and pairwise cosine distances calculated to build a dendrogram using *ward* method.

The TAM and GW neighborhood graphs were built using the previously calculated enrichment scores. To build the graph, we used each m-state in each pair as nodes and connected nodes whose enrichment score was above 8.5 to enrich for highly enriched interacting m-states. We used NetworkX (https://networkx.org) *spring_layout* to facilitate graph visualization.

### Neighborhood in situ enrichment

Neighborhood *in situ* interaction score was built based on the neighborhood enrichment scores. First, m-cells were binned in a hexagonal grid. Next, a vector field was computed by calculating the average enrichment interaction vector between the central hexagon and its neighboring hexagons. Each enrichment interaction vector was calculated as the original distance vector between each hexagon pair multiplied by the total number of m-state pairs found across hexagons multiplied by their enrichment score. The final enrichment *in situ* score was obtained by assigning m-cells to the value of their closest vector module.

### Single m-cell analysis

To analyze tumor m-states, m-cells were pooled from all samples. Cells with a total number of molecules below and above the 5^th^ and 95^th^ quantiles were removed. Top 500 highly variable genes were selected for downstream processing. We normalized the total number of molecules to l2-norm, performed log transformation and *minmax* scaling per each individual sample to remove batch effects. PCA was run and sample specific batch effects were corrected using Harmony. UMAP dimensionality reduction was computed. Gliosarcoma and myeloid cell pools did reveal significant batch effects; in these cases, total counts were normalized, and log-transformed, sample min-max scaling and Harmony correction were not required. Clustering was not necessary as m-states were used for further analysis.

### Data visualization

We used FISHscale (https://github.com/linnarsson-lab/FISHscale) for the spatial visualization of RNA molecules and FISHspace (https://github.com/linnarsson-lab/FISHspace) for the visualization of m-cells as polygons. FISHspace software was developed based on the Python package stlearn^78^, with some added functionalities for visualization and pseudotime analysis.

## DATA AVAILABILITY

EEL-FISH molecule coordinates, accompanying H&E images and Xenium data are available at BioImage Archive. Raw EEL-FISH image data will be shared by the lead contact upon request. Corresponding gene expression data have been deposited on google storage and can be accessed at https://github.com/linnarsson-lab/glioblastoma_wound_response. The m-cell segmented data is available in AnnData format compatible with scanpy, squidpy, and FISHspace. The EEL-FISH RNA coordinates are available as parquet files compatible with FISHscale. Raw scRNA-seq data are available at European Genome-phenome Archive under Accession number TBA. The dataset can also be browsed and downloaded at CELLxGENE (https://cellxgene.cziscience.com/collections/113a558a-e96e-4643-81db-140e95c58578 ).

## CODE AVAILABILITY

All original code has been deposited at https://github.com/linnarsson-lab/FISHscale for the analysis and visualization of RNA molecules and https://github.com/linnarsson-lab/FISHspace for m-cell analysis and visualization. Single-cell RNA sequencing data were analyzed using Cytograph-Shoji (https://github.com/linnarsson-lab/cytograph-shoji). Code for general data visualization can be found at https://github.com/linnarsson-lab/glioblastoma_wound_response.

## ACKNOWLEDGEMENTS

We thank Anna Johnsson, Karolinska Institutet, for grant management and all members of the Linnarsson lab for helpful discussions. We thank the Biomedicum Imaging Core Facility, Karolinska Institutet, and the In Situ Sequencing Infrastructure Unit, SciLifeLab Stockholm, for their services. In addition to author O.P., we would like to thank neurosurgeons Margret Jensdottir and Jiri Bartek and all other neurosurgeons and nurses at the Department of Neurosurgery at Karolinska University Hospital for tissue collection, and patients for donating tissue. This work was supported by Region Stockholm (support to J.K.J.) and the Erling-Persson Family Foundation (Atlas of childhood disease), Hjärnfonden (FO2023-0309), the Swedish Research Council (2022-01248), and the Torsten Söderberg Foundation grants to S.L.

## AUTHOR CONTRIBUTIONS

O.P., J.K.J., and S.L. conceived and designed the multi-region scRNA-seq study. J.K.J. and L.H. performed scRNA-seq experiments. J.K.J., K.S. and S.L performed scRNA-seq data analysis. S.L., A.M.A., and J.J. conceived the spatial transcriptomics study. J.J. and A.M.A. performed EEL-FISH experiments, I.K. performed other stainings. R.K. derived organoids from primary tumor tissue. R.K., M.C.M.F. and I.K. performed organoid experiments. L.E.B., A.M.A. and J.J. optimized the EEL-FISH protocol. A.M.A. performed data analysis. S.C. developed and performed image analysis. E.K. and J.B.M. performed flow cytometry experiments and analysis. O.P. and J.K.J. provided tumor specimens and clinical annotation. A.S. performed pathology analysis. A.M.A., J.J., J.K.J. and S.L., interpreted the results and wrote the manuscript. All authors provided critical feedback and helped shape the project, analysis and the manuscript.

## COMPETING INTEREST DECLARATION

S.L. is a paid scientific advisor to Moleculent and a shareholder in EEL Transcriptomics AB, which owns the intellectual property to the EEL method (“Spatial RNA localization method”, provisional US patent application 63/139,701). All other authors declare that they have no competing interests.

## ADDITIONAL INFORMATION

Supplementary Information is available for this paper. Correspondence and requests for materials should be addressed to the lead contact, Sten Linnarsson.

## EXTENDED DATA

**Extended Data Figure 1.**
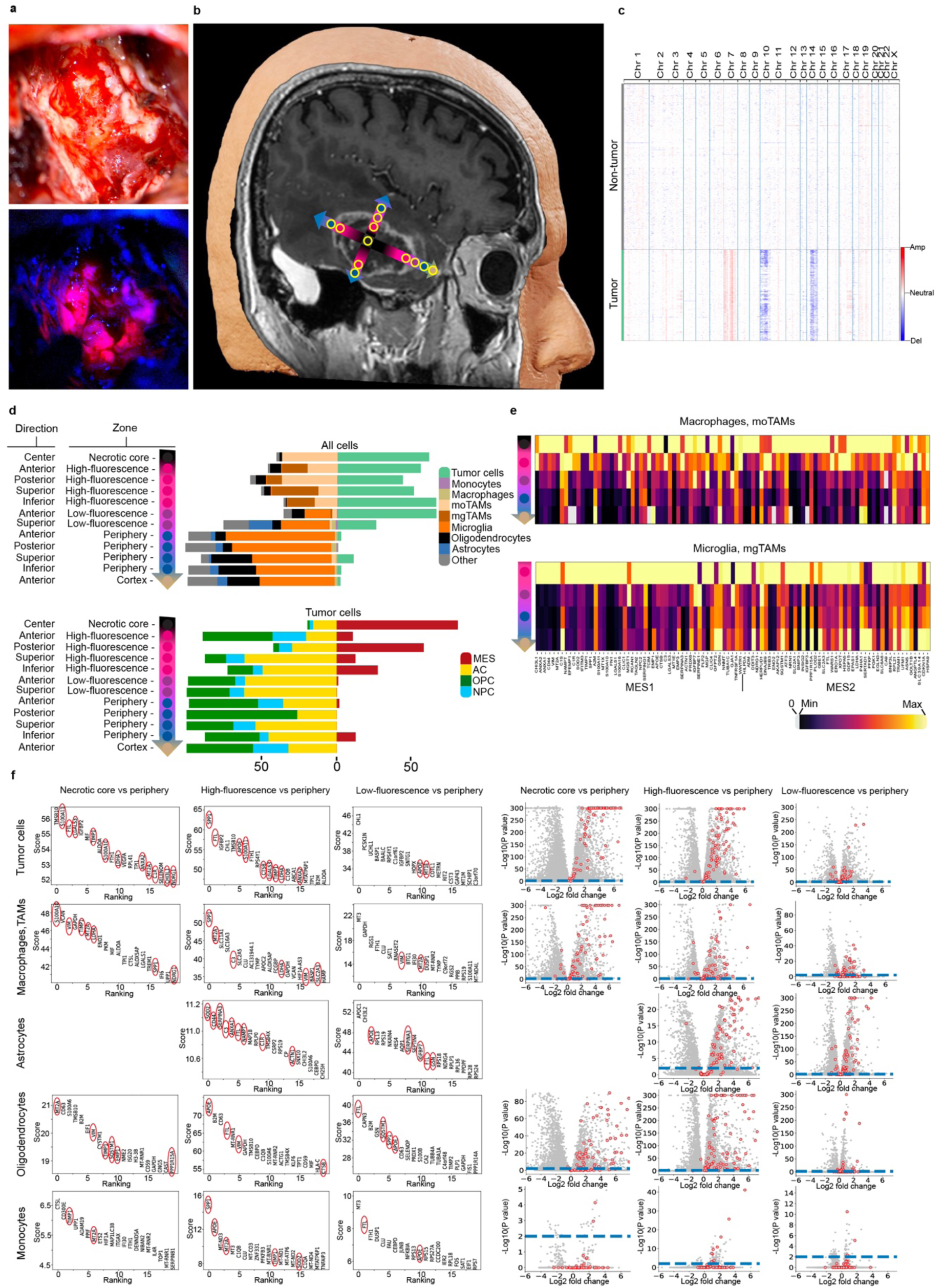
scRNA-seq analysis of multi-region sample SL040. **a**, Images of GBM surgery. Top: Surgical area visualized in white light. Bottom: Same area with fluorescence microscopy in blue light visualizing areas with high-fluorescence (red), low-fluorescence (pink/purple) and no fluorescence (blue and black). **b**, 3D reconstruction of MRI in Brainlab Elements, sagittal plane of SL040. Arrows are colored according to fluorescence and show directions of sampling for scRNA-seq. Yellow circles schematically visualize sampling locations. **c**, Heatmap of the inferred CNV profiles in SL040 across chromosomes (columns) and single cells (rows) from scRNA-seq data. Annotation of cell types (tumor, non-tumor) based on gene expression profiles and clustering on the tSNE. **d**, Top: Bar charts of scRNA-seq cell type proportions in all 12 sampling locations annotated by direction and zone. Bottom: Bar charts of tumor cell state fractions in sampling locations. **e**, MES-like gene expression in the macrophage lineage (macrophages, moTAM) and microglia lineage (microglia, mgTAM) across zones (no microglia lineage cells in necrotic core). The colormap shows linear gene expression from the minimum to the maximum value per gene, with zeros shown in gray. **f**, Differential gene expression by zone (columns) and cell type (rows). No analysis of astrocytes in the necrotic core due to small sample size (number of cells = 2). Left: ranking of the top 20 differentially expressed genes between tumor zones by z-score. MES-like genes are encircled in red. Right: volcano-plots of differentially expressed genes between tumor zones, MES-genes are encircled in red (blue line, q < 10^−2^).

**Extended Data Figure 2.**
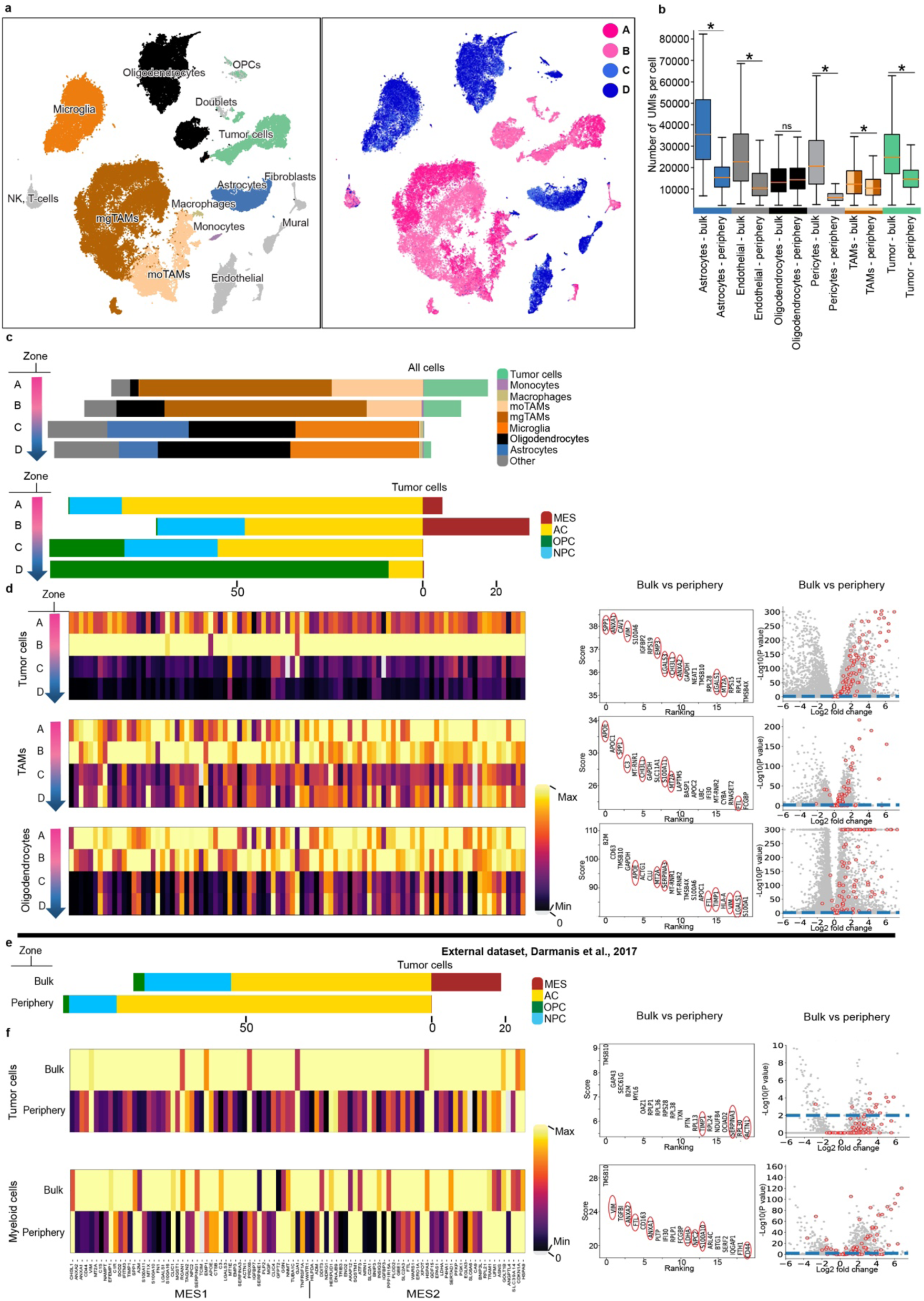
scRNA-seq analysis of multi-region sample SL057 and an external dataset. **a**, t-SNE representation of scRNA-seq data from sample SL057, spanning from tumor bulk (A, B) to periphery (C, D). **b**, Distribution of total UMIs per cell in various cell types by sampling location (bulk/periphery) in SL040 and SL057. Boxes are centered on the median and represents the first to third quartiles. Whiskers extend 1.5 interquartile range above and below the boxes. Asterisks indicate significant differences (Bonferroni-adjusted Mann-Whitney, one-sided (“greater”) and p < 1e10^−18^) for all cell types except oligodendrocytes (p > 0.05). **c**, SL057. Top: bar charts of scRNA-seq cell type proportions across all sampling locations. Bottom: bar charts of tumor cell state fraction across all sampling locations. **d**, Left: MES-like gene expression across cell types and zones in SL057. The colormap shows linear gene expression from the minimum to the maximum value per gene, with zeros shown in gray. Right: Differential gene expression by cell type (rows) and zone (columns). First column from left: ranking of the top 20 differentially expressed genes by z-score. MES-genes are encircled in red. Second column from left: Volcano-plots, MES-genes are encircled in red (blue line, q < 10^−2^). **e**, **f**, External multi-region sample dataset from four patients^27^. **e**, Bar charts of tumor cell state fractions in sampling locations (bulk and periphery). **f**, Left: MES-like gene expression signature, Right: differential gene expression across cell types and zones.

**Extended Data Figure 3.**
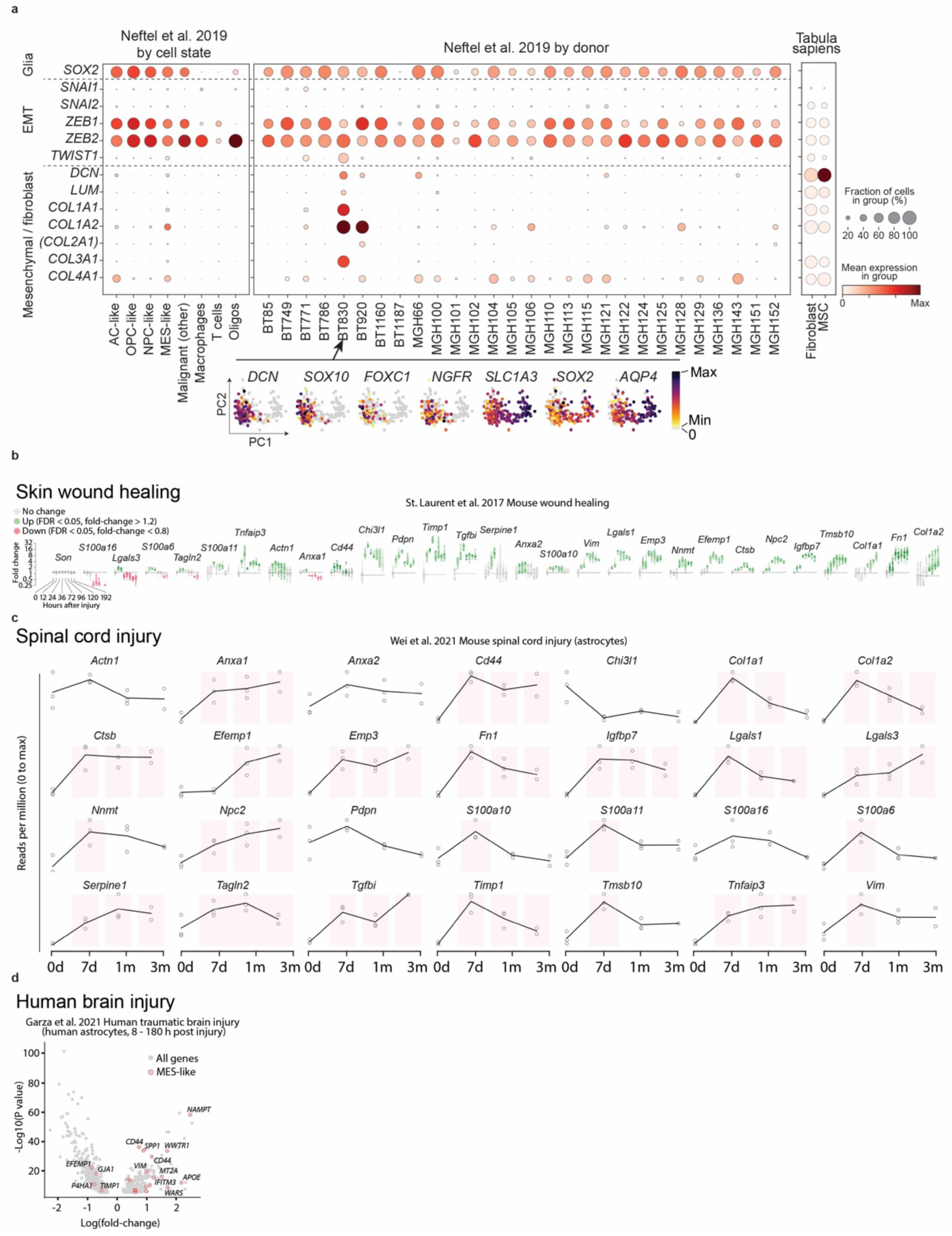
Mesenchymal gene expression in MES-like cells and wound healing. **a**, Expression of mesenchymal marker genes in published datasets, Neftel *et al.*^5^ and Tabula sapiens^48^. Mean expression is not necessarily comparable between datasets. Parentheses around *COL2A1* indicate that this gene is expected to be absent in mesenchymal cells. Arrow: putative gliosarcoma tumor. EMT, epithelial to mesenchymal transition. MSC, mesenchymal stem cell. At bottom, PCA projections of the cells from sample BT830, showing the expression of selected genes. Zeros are plotted in grey. **b**, Regulation of MES-core^37^ genes during mouse skin wound healing^36^. Each plot shows the time-resolved expression of a MES-core gene (*SON* is shown as an example of an unregulated non-MES gene). Green, significantly up-regulated (FDR < 0.05, fold-change > 1.2); red, significantly down-regulated (FDR < 0.05, fold-change < 0.8). **c**, Regulation of MES-core genes in mouse spinal cord injury^38^. Red shading indicates significant up-regulation (FDR < 0.05). **d**, Regulation of MES-like^5^ genes in human traumatic brain injury^39^. Red shows significantly regulated MES-like genes (FDR < 0.05; 18 up, 4 down).

**Extended Data Figure 4.**
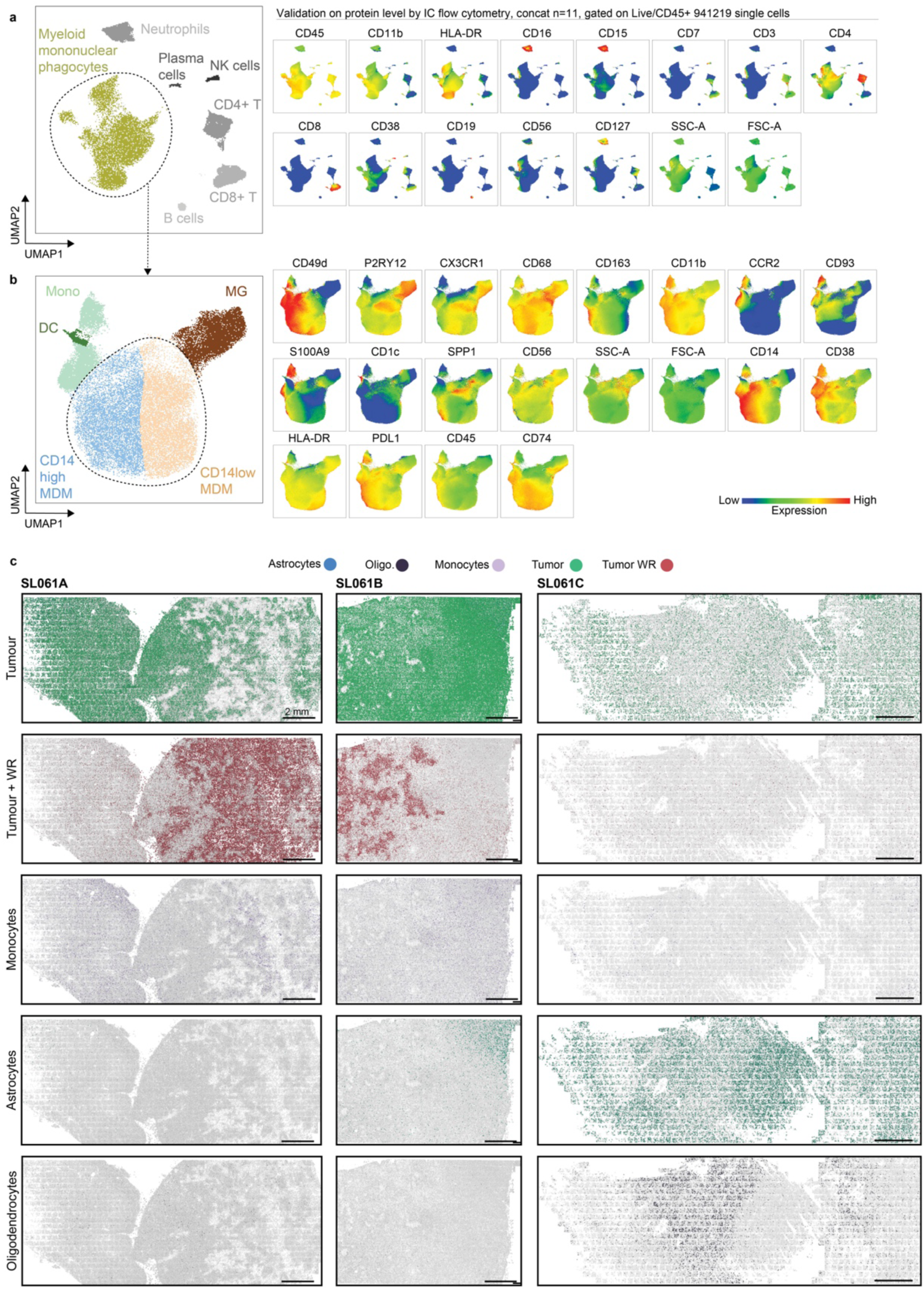
Flow cytometry validation of immune cell populations and non-malignant versus non-malignant cell state classification in EEL-FISH. **a**, Left: UMAP representation of flow cytometry data from concatenated live immune cells (CD45^+^) from 11 GBM samples not included in EEL-FISH cohort with gated myeloid mononuclear phagocytes. Right: corresponding cell identity marker expression. **b**, Left: UMAP representation of myeloid mononuclear phagocytes from A, showing monocytes (Mono), dendritic cells (DC), microglia (MG) and CD14^high^ (blue) and CD14^low^ (orange) monocyte-derived macrophages (MDM). Right: corresponding cell identity marker expression. **c**, Spatial distribution of malignant cells (WR and non-WR), astrocytes, oligodendrocytes and monocytes in a sample with multi-regional representation, SL061, from high-WR (left) to a low-WR (right) region.

**Extended Data Figure 5.**
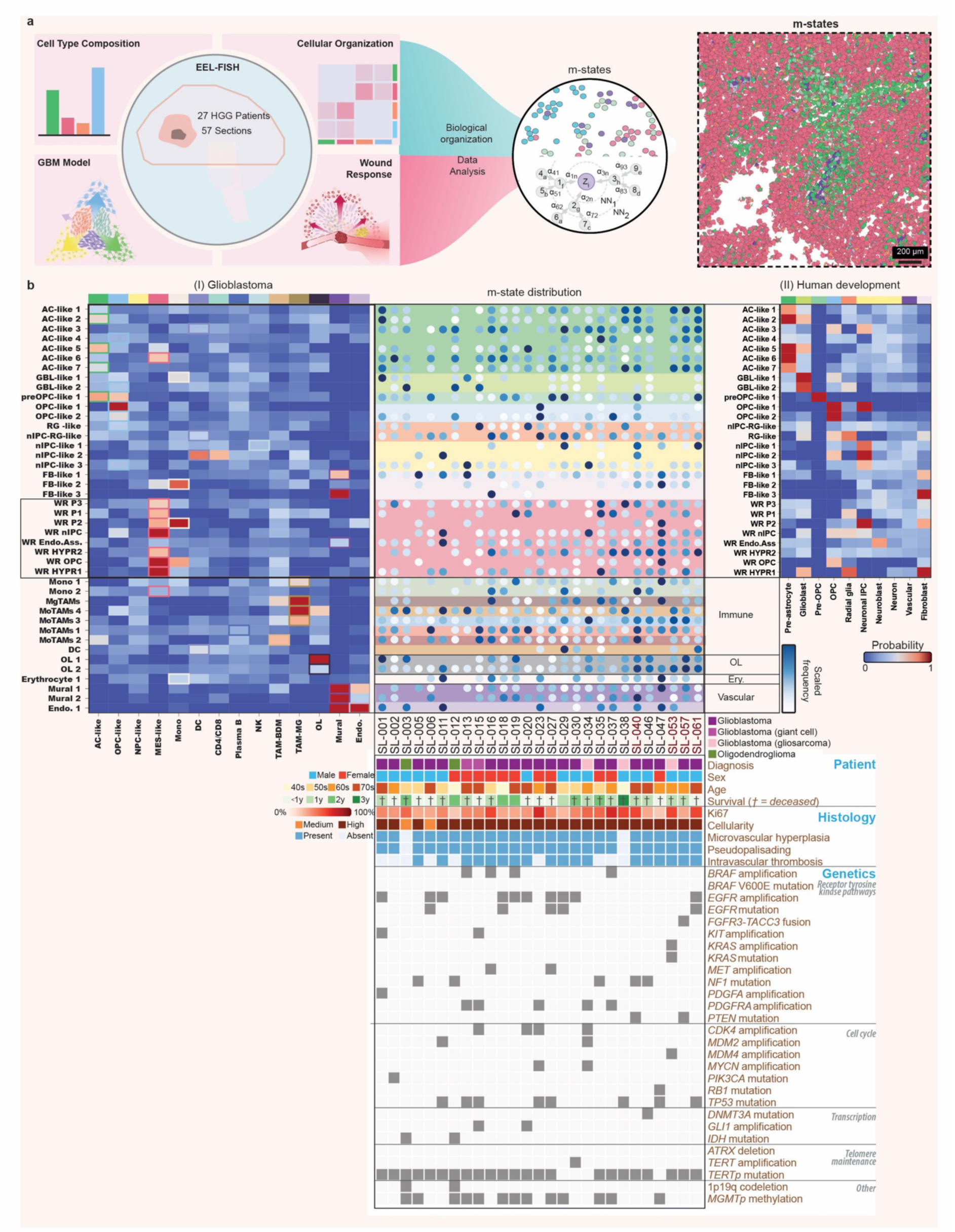
Tangram mapping of m-states, m-state distribution across samples and pathology report summary. **a**, EEL-FISH analysis pipeline schematic for defining common m-states. **b**, Tangram mapping of m-states to scRNA-seq GBM reference^41^ (left-most panel). Tangram mapping of m-states to a scRNA-seq human fetal development reference^42^ (right-most panel). Dot plot of m-state distribution in samples from every patient (middle panel). Pathology report summary, patient information and tumor mutational profiles (bottom-panel). Names of multi-region samples are highlighted in dark red.

**Extended Data Figure 6.**
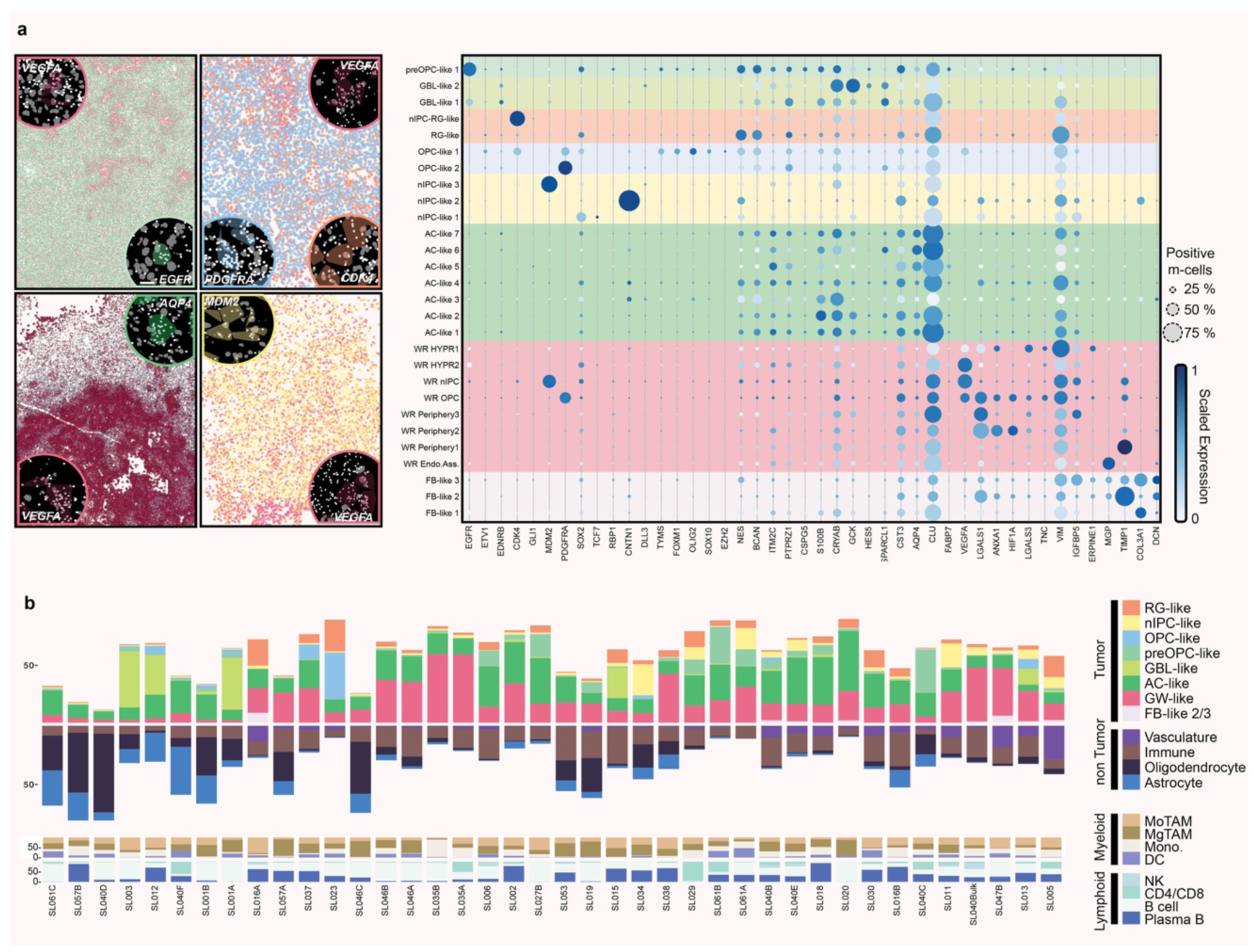
Molecular characterization of GBM m-states using EEL-FISH. **a**, Left: WR marker VEGFA expression in samples with different predominant glia-like m-states (top left quadrant – sample SL061B, top right – SL023, bottom left – SL035B, bottom right – SL034). Colored polygons represent m-cells, gray polygons represent detected cell nuclei. M-state colors correspond to the right panel. Right: Dot plot showing the scaled marker gene expression for each tumor cell m-state. Dot size indicates the percentage of m-cells expressing each gene. **b**, Top: tumor and non-tumor m-state proportions across samples. Bottom: myeloid and lymphoid compartment composition across samples as represented by Tangram probabilities (percentage) for each scRNA-seq reference myeloid and lymphoid cell type as the sum of Tangram probabilities for all sample m-states.

**Extended Data Figure 7.**
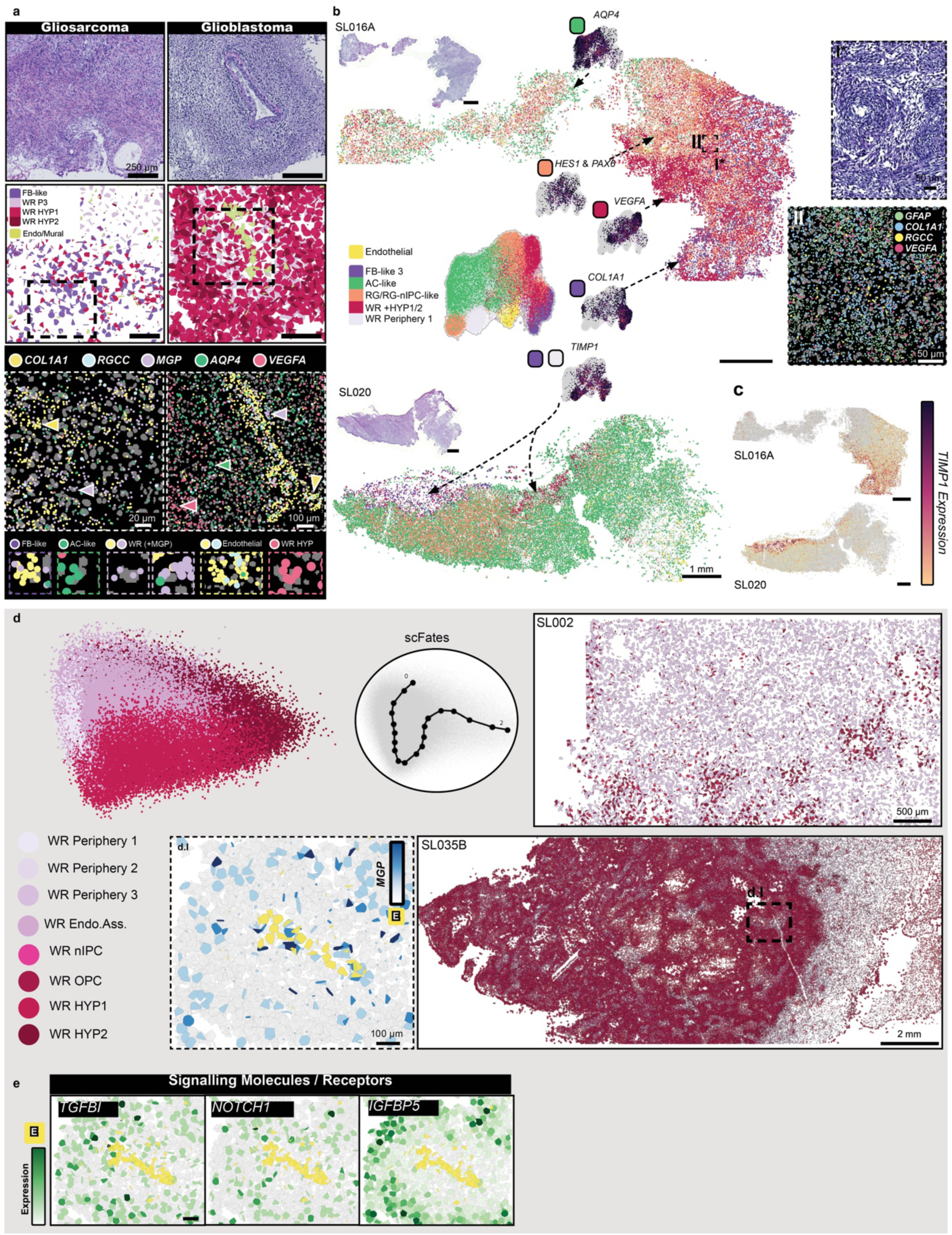
Spatial and molecular characterization of fibroblast-like molecular states and generalized wound response in GBM. **a**, H&E staining, MES-like m-states, and marker gene expression in gliosarcoma (sample SL020) and glioblastoma (sample SL035B). **b**, Spatial distribution of FB-like and WR m-states in gliosarcoma. UMAP representation of endothelial, FB-like, WR and non-WR m-states in gliosarcoma. **c**, *TIMP1* expression in FB-like areas in gliosarcoma. **d**, Top left: Inferred pseudotime trajectory of WR states across all GBM samples. Spatial distribution of WR states in samples SL002 and SL035B. d.I: *MGP* expression in WR m-cells in a perivascular niche in sample SL035B. **e**, Expression of *TGFBI*, *NOTCH1* and *IGFBP5* in WR m-cells in a perivascular niche in sample SL035B (same area as d.I).

**Extended Data Figure 8.**
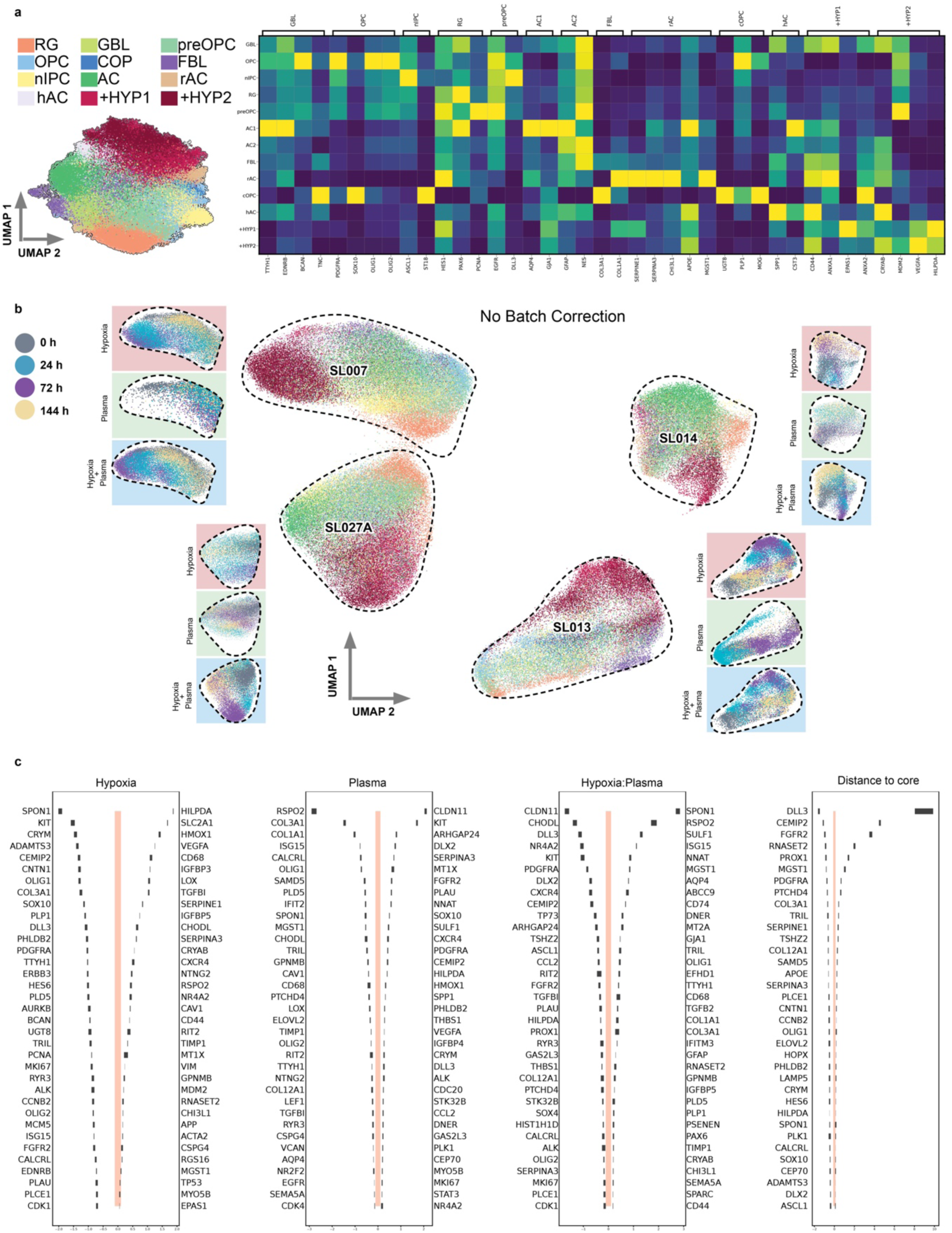
Time-resolved changes in glioblastoma organoid molecular states under hypoxia, blood plasma or both. **a**, Left: UMAP representation of Xenium GBO data, batch corrected by patient. Right: heatmap of marker genes for each m-state. **b**, UMAP representation of non-batch corrected Xenium data across different culturing conditions and time points. **c**, Posterior distribution (quantile 5% and 95%) for the effect size of each gene under hypoxia, blood plasma, both and by distance to organoid core.

**Extended Data Figure 9.**
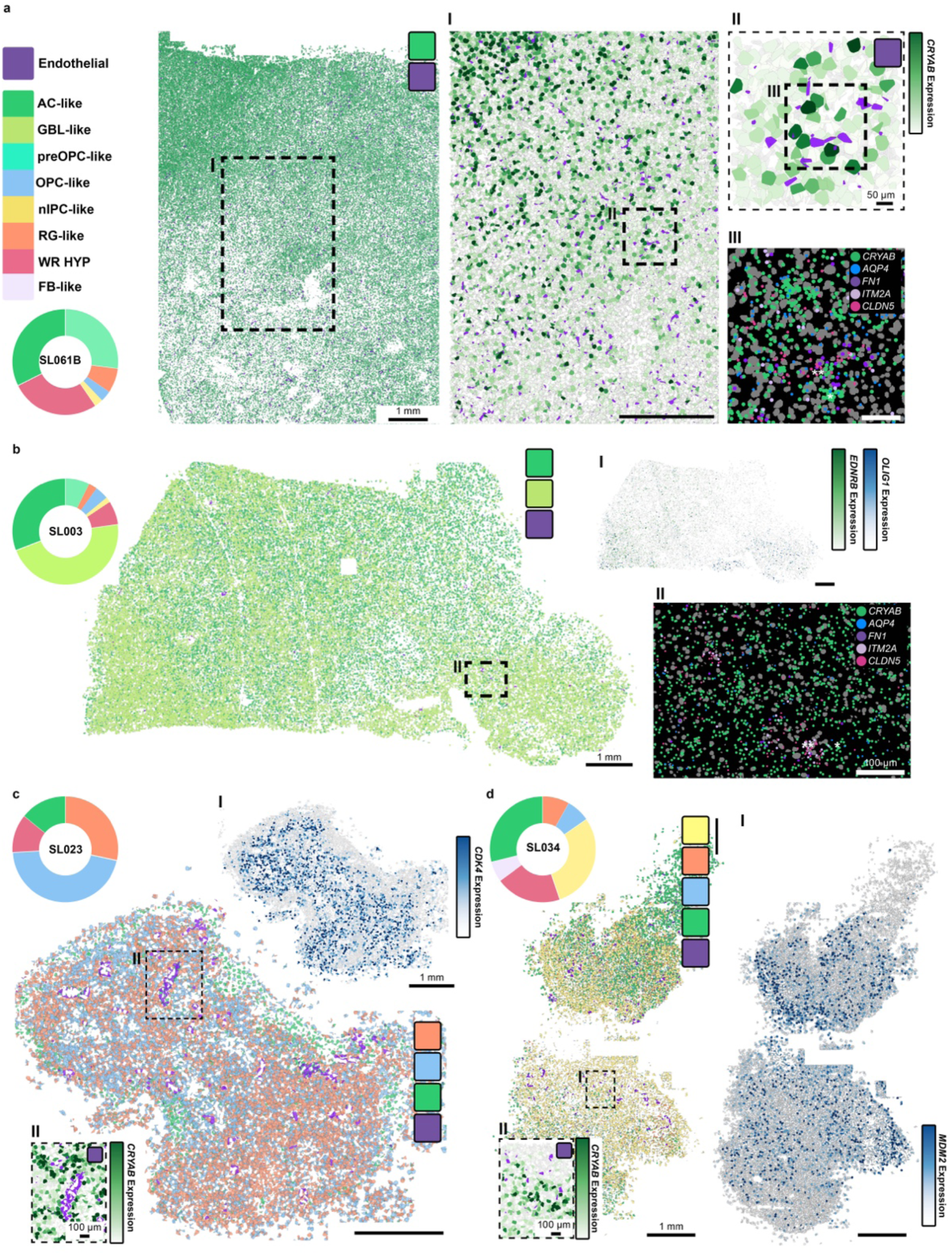
AC-like and endothelial m-cell co-localization in GBM and oligodendroglioma. **a**, Spatial distribution of AC-like and endothelial m-states in sample SL035B; pie chart shows m-state proportions. I: Co-localization of *CRYAB* expression and endothelial m-cells. II: Close-up view of region I. III: Co-expression of AC-like markers *CRYAB* and *AQP4* (*) near endothelial cell marker genes *FN1*, *CLDN5* and *ITM2A* (**). Gray polygons represent m-cells with zero expression. **b**, Spatial distribution of AC-like, GBL-like and endothelial m-cells in oligodendroglioma sample SL003; pie chart shows m-state proportions. I: Expression of *OLIG1* and *EDNRB* in sample SL035B. Gray polygons represent m-cells with zero expression. II: Co-expression of AC-like markers *CRYAB* and *AQP4* (*) near endothelial cell marker genes *FN1*, *CLDN5* and *ITM2A* (**). **c**, Spatial distribution of OPC-, RG-, AC-like and endothelial m-cells in sample SL023 with corresponding *CDK4* expression. Gray polygons represent m-cells with zero expression. I: *CRYAB* expression surrounding endothelial m-cells. **d**, Spatial distribution of nIPC-, RG-, OPC-, AC-like and endothelial m-cells in sample SL034 with corresponding *MDM2* expression; pie chart shows m-state proportions. Gray polygons represent m-cells with zero expression. II: *CRYAB* expression surrounding endothelial m-cells.

**Extended Data Figure 10.**
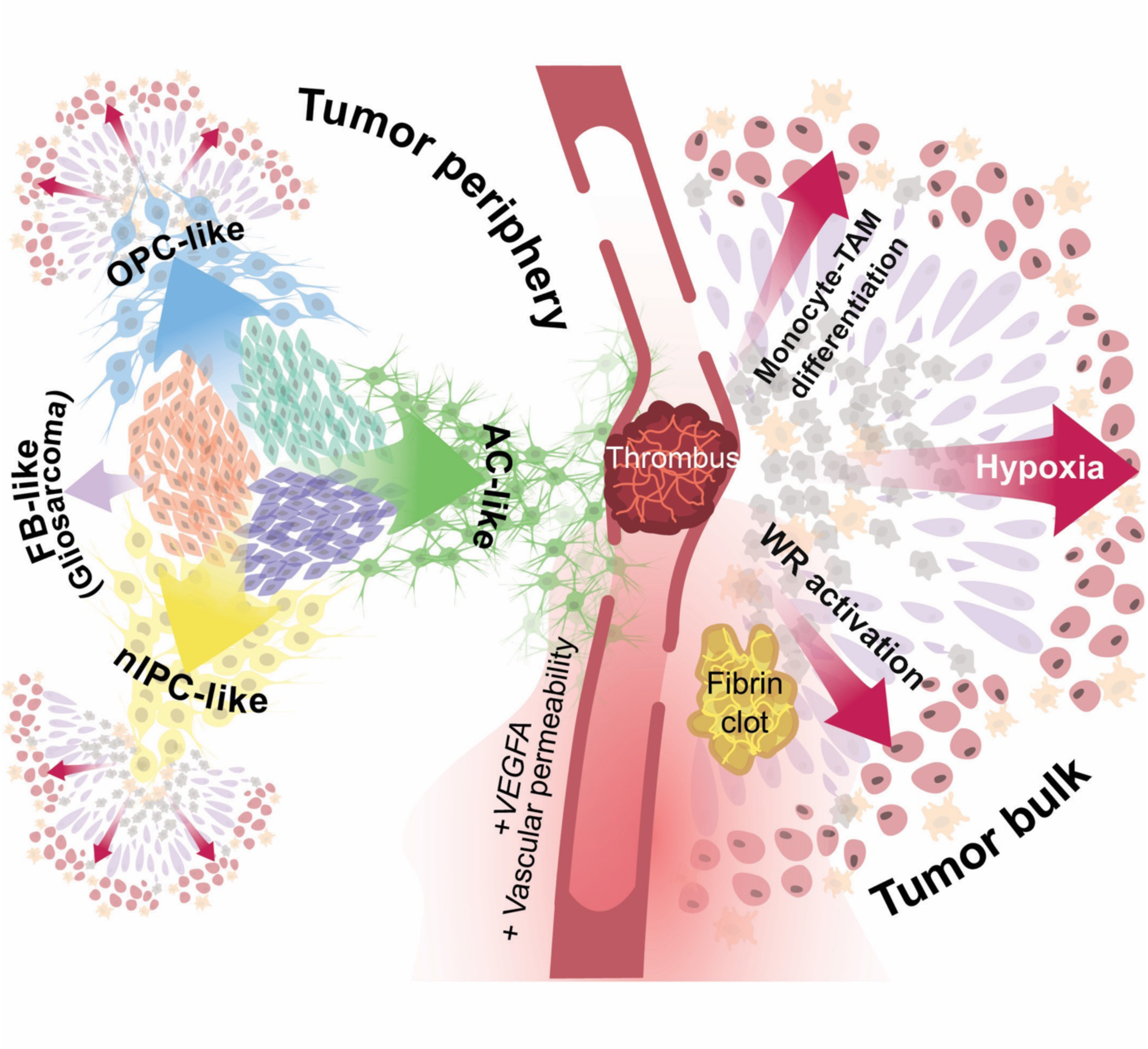
Model of GBM organization by wound response. Glia-like cell states proliferate and drive tumor growth. Tumor bulk formation brings about tissue disruption, vascular dysfunction, hypoxia, promoting hemostatic responses and monocyte infiltration. Consequently, WR states are activated, monocytes are sequestered and differentiate into TAMs, perpetuating the futile wound response.

## Notes

https://github.com/linnarsson-lab/glioblastoma_spatial/

### Summary of Updates

This version of the manuscript combines previous EEL-FISH data from a large patient cohort with deep scRNA-seq on two of multi-region GBM cases, and Xenium Spatial transcriptomics on GBM organoids.

